# Reversible and reusable compartmentalized thermoplastic chip for coculture of dorsal root ganglion neurons

**DOI:** 10.1101/2024.12.10.627139

**Authors:** Solène Moreau, Raul Flores-Berdines, Anne Simon, Tatiana El Jalkh, Guillaume Taret, Anna Fomina, Céline Dargenet-Becker, André Estevez-Torres, Sophie Bernard, Hugo Salmon

## Abstract

Compartmentalized microfluidic chips play an important role in understanding the cellular mechanisms involved in neurodegenerative disorders. Dorsal root ganglia are a well-established model for modelling the peripheral nervous system (PNS), but their development on a chip remains limited. Furthermore, it would be beneficial for the devices to be openable in order to access the biological material inside for analyses. Easy to prototype and biocompatible, styrenic block copolymers (SBC) are an alternative to polydimethylsiloxane (PDMS) that offer both reversible and permanent bonding properties. This paper presents a fast and straightforward method to produce compartmentalized SBC chips. The study validates the culture of murine dorsal root ganglia explants, comparing it to the standard methods, to obtain a model of the PNS. Moreover, the reversible bonding properties of the SBC permit the reuse of the chip with a quick and easy cleaning protocol. It provides direct access to the cells, opening the way for imaging and molecular biology analysis. The comparison of the resources required to produce PDMS and SBC chips highlights the importance of moving to reusable devices. These detachable, easy-to-manufacture and sustainable all-thermoplastic platforms provide an alternative way of prototyping compartmentalized devices *for in vitro* PNS modeling.

## 1. Introduction

The reduction of animal numbers and their replacement by alternative experimental methods are two of the 4R principles (reduction, replacement, refinement, and responsibility) and constitute a major strategic ethical challenge, prompting the development of *in vitro* models that can be used instead of *in vivo* studies involving animal models. Despite the rise of microphysiological systems, in neuropharmacology, the complexity of the physiological processes and the long time scales might explain why only few neurological conditions have been modeled to date.^1^ While industrial sectors have played a role in democratizing these tools, reducing their demand on laboratory resources and broadening their applications in neurobiology could further advance this effort.^2^

Microfluidic devices for compartmentalized culture have offered the possibility to study the neuronal functions and structures in a highly controlled way.^3,4^ Robert Campenot was the first to develop a device that enabled local control of axons and somas.^5^ The aim was to physically separate the cell bodies from the axons to mimic the diverse microenvironments that a neuron might encounter. Subsequently, a variety of microfluidic devices have been developed for neuronal studies. Taylor initiated a two-compartment microfluidic device connected by straight microchannels ^6^ for the study of axonal injury, regeneration and transport. Samson *et al.* employed a five-parallel culture chamber configuration to circumvent cross-contamination and to establish an environmentally isolated neuronal population with synaptically interconnected cells.^7^ Courte *et al*. designed a microchip with arc-shaped microchannels to control the orientation and filtering of axonal growth.^8^ Sharma *et al.* proposed a keyhole design to model human nerves,^9^ while Park *et al.* developed central circular chamber designs to promote axon growth in microchannels and the formation of a denser axonal network layer.^10,11^ Electrospun aligned nanofiber-integrated compartmentalized microfluidic neuron culture systems have been developed by Luo *et al.* to improve axon alignment, length and stability.^12^ Two-dimensional (2D) and three-dimensional (3D) compartmentalized co-culture systems on microdevices have been developed to analyze the interactions between neurons and glial cells in both the central nervous system (CNS)^11,13^ and in the peripheral nervous system (PNS).^9,11,14–16^

Most of the PNS models focus on motor neurons, while there are only a few studies on sensory neurons on chip.^17–20^ However, these studies focus on co-cultures of sensory neurons and oligodendrocytes, a glial cell that is specific to the central nervous system. Dorsal root ganglia (DRGs) represent a well-established model for PNS modeling, but their development on a chip remains limited. They mainly focus on pain signaling^21^ and therapeutics^22^ as well as on “nerve-on-chip”.^23,24^ In 2023, Mutschler *et al*.^25^ proposed the first co-culture of neurons from dissociated DRGs and Schwann cells on a compartmentalized microfluidic device. They highlighted the challenge of culturing DRGs as explants in microfluidic chips. Notably, Park *et al.* were the only group to successfully culture a chick DRG explant in a compartmentalized microfluidic device.^26^ To facilitate the seeding of the explant, they designed a device with an open-top chamber.

Compartmentalized systems and organ-on-chip devices are becoming more widespread in the field of Biology, requiring a re-evaluation of their sustainability and of the materials used in their manufacture for specific applications. Currently, polydimethylsiloxane (PDMS) is predominantly used due to its high oxygen permeability, while polystyrene (PS) is favored for large-scale production and alignment with biological standards.^27^ However, there are no neurobiological applications that uses materials combining the advantages of PDMS and PS, such as styrenic block copolymers (SBCs).^28^ Reversible bonding of SBCs would be an advantage for accessing tissues during culture and for reusing devices.^29–31^ Moreover, as the organ-on-chip industry is expanding rapidly and access and production costs are becoming critical for laboratory resources and carbon footprint,^32^ there is no known comprehensive assessment or methodology to evaluate the resources involved.

Here, we describe a fast and straightforward method for the fabrication of compartmentalized SBC chips and demonstrate their reversible bonding and reusability for the culture of compartmentalized peripheral nervous system model. We first propose a simple process for fabricating SBC chips with an axisymmetric design adapted to anisotropic compartmentalized culture, particularly suitable for primary DRG explants. We confirm the theoretical compatibility of the design with neuron culture using analytical and finite element models. The viability and axonal growth of *E*13.5 mouse embryonic DRGs within SBC and equivalent PDMS chips were then compared. We present a rapid and reliable cleaning procedure for SBC chips and evaluate their reusability. We also demonstrate the potential of reversible bonding chips for direct imaging using electron microscopy, as well as for biological analysis techniques (immunostaining, Western blot, RT-qPCR). Finally, we discuss the resources (energy, time, cost and CO_2_ equivalent emission) involved in the production of a PDMS compared to an SBC chip.

## 2. Results & discussion

### 2.1. Development of a compartmentalized SBC chip for coculture of DRG neurons

We developed a design and a workflow to produce axisymmetric compartmentalized 6-plexed SBC devices. With the exception of the photolithography step, which is necessary for wide channels, the fabrication process of SBC microfluidic chips **(Figure 1A)** is an out-of-the-cleanroom process. The first phase was to replicate the SU-8 mold in an epoxy mold, which is more compatible with hot embossing. It was used for 2 years, embossing more than 200 chips without any visible damage. The replication of the molds was confirmed by profilometry (**Figure SI.1**). These open-top devices are made of two layers that are simple to assemble thanks to the adhesive properties of SBC. A first 1 *mm*-thick layer containing the structures and a 2 *mm*-thick reservoir layer to increase the volume capacity and prevent rapid evaporation of the culture media (**Movie SI.1**) as well as shear stress during pipetting. The central chamber without the reservoir can only hold 7 µL, whereas it reaches a volume of 63 µ*L* when the second layer is added. Due to the radial growth of axons in the DRGs, we designed a circular central chamber with a diameter of 4 *mm* to facilitate the penetration of axons into the microchannels **(Figure 1B)**. The dimensions of the microchannels have been chosen to enable compartmentalization of somas in the central chamber and axons in the peripheral chambers.^6,33^ Treatments can be added in the outer chambers without flowing back to the central chamber. The high fluidic resistance of the microchannels creates a small but sustained flow between the chambers, thereby counteracting the diffusion (**Figure SI.3**). This fluidic isolation allowed 6 independent peripheral chambers to be created. The sTPE devices bond easily to polystyrene (PS) **(Figure 1C)**, making them adaptable to already existing basic culture plates. The Young’s modulus of the SBC polymer is 1.15 *MPa*^28^ which is in the same range as PDMS (0.5 − 3 *MPa*). In addition, this material is biocompatible, gas permeable, optically transparent and exhibits reversible bonding that can withstand 1 *bar* even after 10 sealing/unsealing cycles,^34^ making it a suitable candidate for cell culture. The air plasma hydrophilization remains stable in time for at least one week for the SBC chips. The culture of peripheral neurons was demonstrated using either explants or dissociated embryonic DRGs harvested from *E*13.5 mouse embryos.

**Figure 1:**
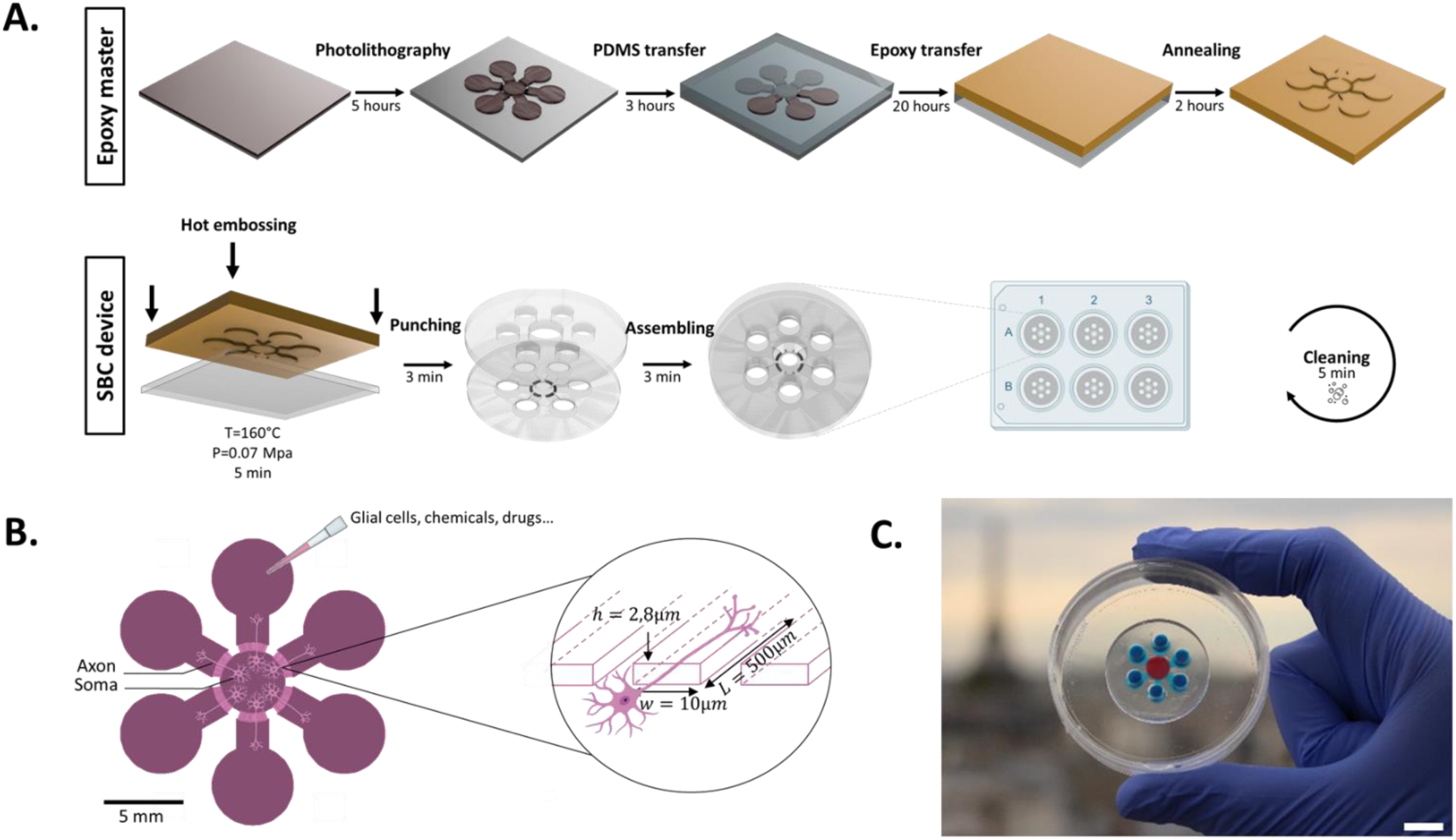
Fabrication and design of the thermoplastic chip. **A.** Fabrication process of the two-layer microfluidic chip in styrenic bloc copolymers (SBC). **B.** Scheme of the design of the chip. Somas are compartmentalized in the central chamber, while axons grow through the 292 microchannels. **C.** Image of the device filled with food dyes. Scale bar = 1 *cm*.

The device was also compatible with dissociated DRGs. It has been described that the radial flow pressure resulting from the circular fluidic design facilitates the positioning of cells in close proximity to the microchannels inlet.^10,11,35^ Moreover, the circular design allows for axon coverage that is more than 3.5 times higher than that of square designs.^10^ It allows the cultivation of both dissociated DRGs and explants while most of the DRGs cultured on chips are dissociated. Overall, this model provides an experimental framework to study different aspects of Schwann cell development such as proliferation, migration, differentiation, and myelination of axons.^36^ These co-cultures of neurons, Schwann cells and fibroblasts were placed directly into the central chambers and cultured within the SBC chips bonded to 6-well PS plates. The cultures could be maintained for one month within the devices, and no leakage or contamination was observed.

### 2.2. A shear stress compatible with DRG culture on a chip

In order to assess the shear stress experienced by the DRG in the chip, we performed a Finite Elements Method (FEM) analysis and particle velocimetry. As demonstrated by Liu *et al.*, it is essential to quantify the shear stress forces generated within the microchannels in order to evaluate the suitability of the device for biological studies. We used the FEM to visualize the speed *v* and the shear rate *σ* in a microchannel cross section **(Figure 2A)**, the two physical quantities being intrinsically linked by *σ* = μ𝛻*v*. The following values were obtained for a pressure gradient of 30 Pa, which represents the theoretical maximum pressure that can be applied within the microfluidic device under study, given the finite volume of the reservoirs. The maximum shear stress inside the cross section is 0.8 dyn. cm^−2^. A pressure gradient of 30 Pa is achieved when the central chamber is full and the peripheral chambers are empty. However, this extreme case never occurs in practice when cells are in the chips. During the media change, the pressure reached a maximum of 25 Pa for a few seconds, due to the presence of a small volume of media within the chambers. Experimental validation was performed by measuring the time required for 0.2 µm microbeads to cross the microchannels **(Figure 2B)**. The experiment was started with a pressure of 25 Pa and stopped after 9 h, when the pressure had reached 0 Pa. The flow rate (*Q*) inside the microchannels was calculated from the velocity of the microbeads and the area of the cross section of a microchannel (*A*): *Q* = *v* × *A*. An exponential regression was fitted to the experimental results, with *v*_*max*_ = 132.5 µ*m*. *s*^−1^ and a time constant of 𝜏 = 3.18 ℎ. It was then possible to derive the shear stress inside the microchannels based on the following equation: 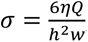, where 𝜂 was fixed as the viscosity of water in Pa.s. For a maximum pressure of 25 Pa, the mean shear stress obtained was 2.73 dyn.cm^-2^.

**Figure 2:**
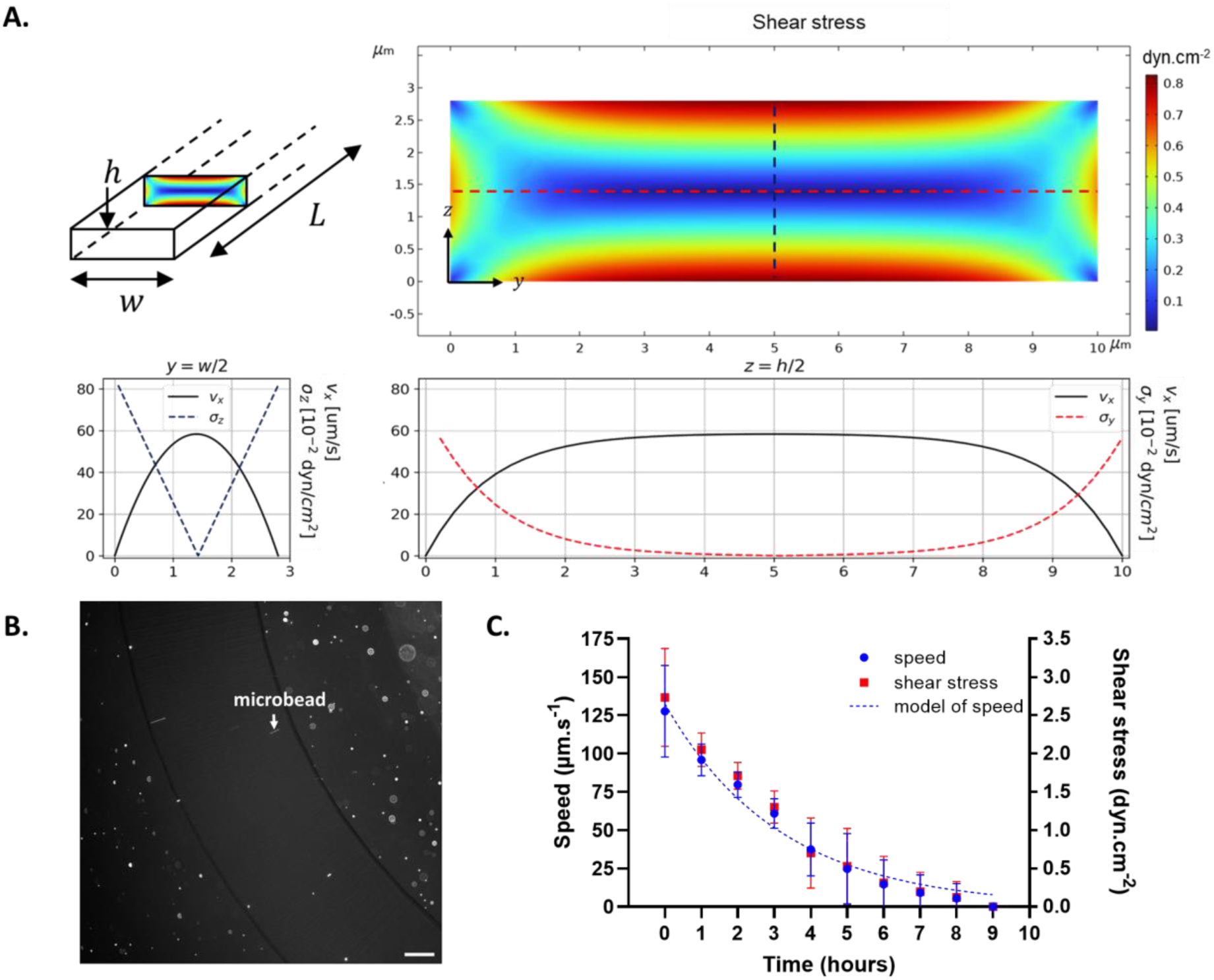
Estimation of the shear stress inside microchannels. **A.** Shear stress field induced by a 30Pa flow over a microchannel cross-section calculated by Finite Element Methods (FEM), with associated analytical plots of the velocity (in black) and shear stress along the y-axis (dashed red line) and along the z-axis (dashed blue line). **B.** Particle image velocimetry in the styrenic bloc copolymers (SBC) chip of 200 nm fluorescent polystyrene particles. Scale bar = 100 μ*m* **C.** Evolution of the flow average speed inside the microchannels (blue curve), fitted exponential model for the average flow speed (dotted blue line), and associated shear stress (red curve) over time for initial pressure conditions of 25 *Pa*.

### 2.3. Viability and axonal growth of DRG culture on a chip

In order to validate the DRG culture on our devices, we assessed the viability and the axonal growth over time and according to the material. For viability, the dissociated cells formed a rather uniform distribution of cells, thereby facilitating the Live/Dead staining. For axonal growth, the axisymmetric growth of the explant is directional, thereby facilitating the segmentation and the quantification of the network. DRGs at E13.5 are at a stage at which we can obtain a high yield of sensory neurons and Schwann cells. Both exhibit high viability and plasticity.^36^ After the DRGs were picked from the extracted spinal cord, dissociated and seeded in PDMS and SBC chips, or in well plates as a control. Their viability was assessed at day *in vitro* (*DIV*) *DIV*3 and *DIV*7. Comprehensive tiles of each culture enable exhaustive quantification and better visualization **(Figure 3A**). All the cells were segmented and counted exhaustively (**Figure SI.6**). At *DIV*3, the cell density was below the expected seeding cell density due to the low survival rate of the primary cells during the seeding step. The number of live cells was significantly higher in the control than in the chips (**Figure 3B**), with more than 1800 cells.mm^-2^ in the 96-well plate against 600 cells.mm^-2^ in the PDMS chips, 880 cells.mm^-2^ in the SBC chips and 930 cells.mm^-2^ in the reused SBC chips at DIV7. However, there was no significant difference in the number of dead cells between all the conditions. The overall number of cells increased by 13 % in the control group between *DIV*3 and *DIV*7, compared to 64% in the PDMS chips. This rate was found to be similar between SBC and reused SBC chips, at 31 % and 35 % respectively. Similarly, DRG explants were fixed at *DIV*3, before the axons reached the microchannels, and at *DIV*7, after the axons had crossed the microchannels **(Figure 3C)**. The growth speed was deduced from the axonal length (**Figure SI.7**). Due to the fluidic isolation, the antibodies were unable to diffuse from the peripheral chambers into the microchannels. As the axons started to cross the microchannels around *DIV*4, it was necessary to detach the SBC chips at *DIV*7 before staining the cells to fully observe them in the peripheral chambers (**Movie SI.2**). However, due to the irreversible bonding of the PDMS, it was not possible to obtain the length of the axon at *DIV*7 inside them. No significant difference was observed between the axonal length in 4-well control and the axonal length in the chips **(Figure 3D)**. From DIV0 to DIV3, the axonal growth speed was 443 ± 33 µm. day^−1^ and decreased to a speed of 324 ± 36 µm. day^−1^ from DIV3 to DIV7.

**Figure 3:**
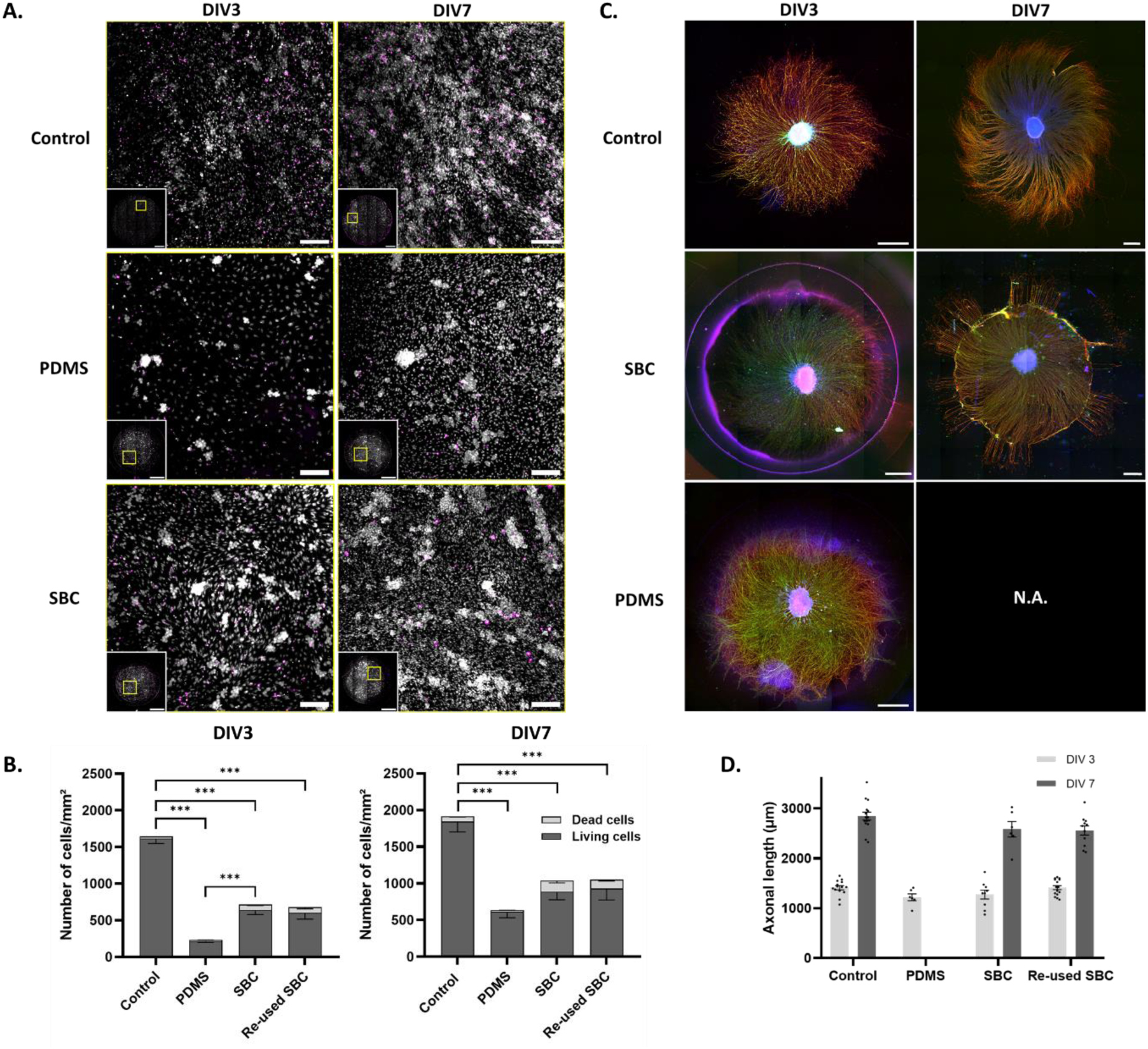
Comparison of viability and axonal growth of Dorsal Root Ganglia (DRG) explants within the chips. **A.** Viability test for a control well from a 96-well plate and a central chamber of a Styrenic Block Copolymer (SBC) and a PolyDiMethylSiloxane (PDMS) chip at *DIV*3 and *DIV*7. The images show all cells (white) and dead cells (magenta). Miniature tiles (bottom left of each image), scale bars = 1.2 *mm*, and its representative frame corresponding to the region in the yellow square, scale bars = 150 µ*m*. **B.** The number of living and dead cells per mm² inside the control wells, PDMS chips, SBC chips and reused SBC chips at *DIV*3 and *DIV*7. DRG 𝑛 = 6 − 24, embryos 𝑒 = 6 − 12. Results are represented as the mean ± SEM. Two-way ANOVA were performed. The alpha value threshold was set at 5% and the *P*-values are represented as follows: ∗ 𝑝 < 0.05, ∗∗ 𝑝 < 0.01, ∗∗∗ 𝑝 < 0.001. **C.** Tiles of DRG explants inside the controls wells, PDMS chips, SBC chips and reused SBC chips at *DIV*3 and *DIV*7. Nuclei (blue), neurofilaments H (green), SMI312 (red), scale bars = 500 µ*m* **D.** Axonal length inside the control wells, PDMS chips, SBC chips and reused SBC chips at *DIV*3 and *DIV*7. No data is available for PDMS chips at DIV7, as it was not possible to measure the axonal length. 𝑛 = 6 − 15, 𝑒 = 6 − 12. Results are represented as the mean ± SEM.

This design enables a uniform treatment in the central chamber and a gradient treatment in the peripheral chambers, due to the high resistance of the microchannels. This feature can be useful for drug treatments; however it prevents antibodies to reach the microchannels during immunostaining constraining us to open the chip to observe the entirety of the axons. In addition, the open-top device facilitates media changes and the CO2 exchange, while minimizing mechanical stress and shear stress compared to a closed-compartment device.^20^ The stable hydrophilization was especially convenient for transport and long-term use compared to PDMS. We compared the maximum shear stress value obtained in our device to the maximum shear stress value that is sustainable for cortical neurons. We considered the value of 5 dyn. cm^−2^which was measured for a 24 h exposure^37^ as a reference, as to our knowledge there are no data on the effect of shear stress on sensory neurons. The maximum experimental shear stress obtained in our device is 1.25 times higher than the maximum theoretical shear stress. However, it is almost 2 times lower than the cell viability limit for cortical neurons. The maximum shear stress in our device is only reached for a few seconds, when the cells are inside, during the change of medium. An experimental study of the effect of the shear stress on the cell viability would help identify the limit in the context of peripheral nervous system (i.e. sensory neurons) but it would require a controlled flow rate. We numerically confirmed that the axonal growth rate is not impacted by the microfluidic chip, assuming a comparable threshold in our biological context. We made tiled images of the central chambers of the chips and of the wells of the controls to confirm the viability of the cells and ensure that our microfluidic architecture does not affect them. The tiles enabled us to monitor viability over time and confirm the homogeneous spatial distribution of dead cells in the chips. In contrast, in a standard well plate, cells tended to accumulate on the walls.

We had to adjust the number of cells seeded for each material to ensure the targeted seeding density remained consistent. Even when using the same 3 mm puncher to open the central chamber of the SBC and PDMS chips, the diameters of the holes obtained are not the same (**Figure SI.2**). This is a consequence of the elongation at break of the SBC (720 %) compared to the PDMS (76 %) one.^27,34^

As the diameters are different, the seeding area is different for both materials. We have a surface area of 7.1 mm^2^ for SBC devices and 3.14 mm^2^ for PDMS ones. However, only the open top parts are different in size, the central chamber is still 4 mm in diameter for both materials, which explains why there are fewer cells per mm^2^in the PDMS devices than in the SBC devices.

The consistently high viability (>80%) across conditions indicates that the cells generally survive well under all tested conditions, which is critical for biological relevance and assay reliability. The axonal growth rate in the chips is comparable to that of the control, indicating that the presence of microchannels has no effect on axonal growth. It is also similar to that observed in the literature, with a rate of around 400µm/day.^38^ The lower cell density in the chips compared to well plates may reflect differences in culture environment, such as nutrient availability or surface interactions, which can influence cell proliferation behavior. The SBC material exhibits denser cell growth at DIV3 and DIV7 in comparison to the PDMS, suggesting that it may offer a better support for cell proliferation. The fact that the reuse of SBC material does not affect viability is encouraging for reproducibility and material stability. In case of in vitro preclinical study, the viability remains relatively high for all three conditions and networks neuron networks develop normally. Further phenotyping would be required to assess advanced structural features such as myelination (Ki67 proliferation marker, c-fos stress marker, MBP and MPZ myelin markers).

### 2.4. SBC chips can be cleaned and re-used up to 3 times

A rapid, simple and reproducible protocol was developed to clean and reuse the chip **(Figure 4A)**. We validated the regeneration of the chip using immunofluorescence, profilometry and electron microscopy. We validated the compatibility of re-used chips using axonal growth assays. The process was completed in less than 3 *m*𝑖𝑛 per device and required only milliQ water, soap, ethanol 70 % and a toothbrush. Chips were sterilized again by UV-sterilization prior to culture. DRGs were grown up to one month in reused chips. It was confirmed that there was no noticeable difference in the surface of the SBC device before and after cleaning. Epifluorescence and SEM images were taken before use, after use and after the cleaning **(Figure 4B)**. Due to the sensitivity of the sTPE to electron beam, cracks appeared during SEM imaging. Using these two microscopy techniques, we confirmed that all the cellular debris present after culture had been removed by the cleaning protocol. After the third cleaning, it was observed that the axons were unable to enter the microchannels **(Figure 4C)**, suggesting that they were obstructed at the entrance. This finding was confirmed by optical profilometry measurement **(Figure 4D)**, which revealed that the microchannels were clogged. The clogging can be readily observed with a brightfield microscope (**Figure SI.9**) and is therefore easily detected. Consequently, each chip can be used up to 3 times before exhibiting irreversible deformation and becoming inoperable. The SBC material also has intrinsic advantages for cell culture compared to the PDMS in our context: the hydrophilization of SBC remains steady, allowing longer storage and reduced risk of cells detachment. The tendency of PDMS to leach and absorb other material can also result in issues when used in conjunction with other materials, particularly 3D printed materials or hydrogels.^39^ The use of SBC materials would facilitate the production of composite devices for more complex applications.

**Figure 4:**
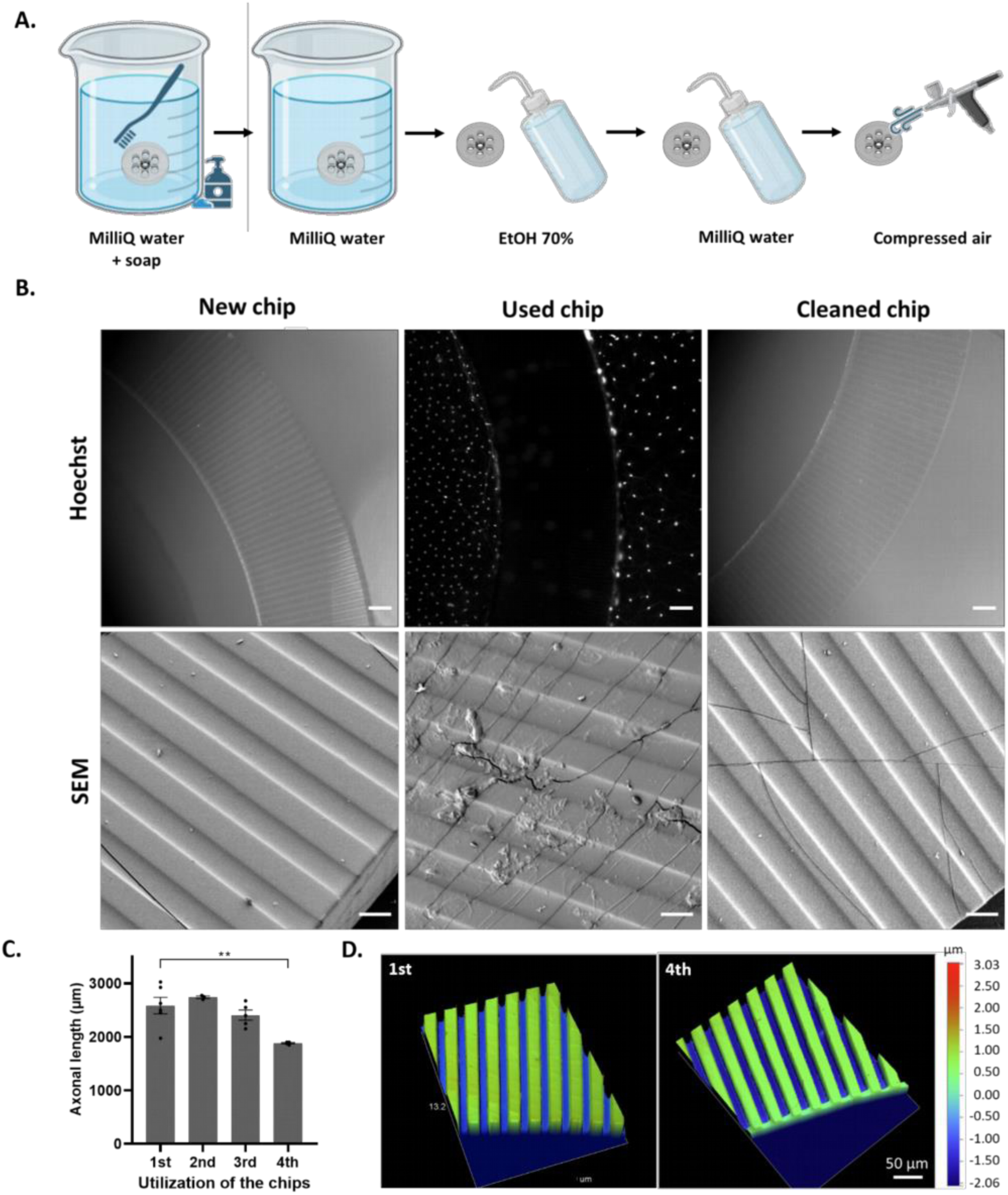
Reusability of Styrenic Block Copolymer (SBC) chips. **A.** Cleaning process of the two-layer microfluidic chip in SBC. **B.** Hoechst staining (scale bar = 100 µ*m*) and SEM (scale bar = 10 µ*m*) imaging on a new, a used and a cleaned SBC chip. **C.** Axonal length of Dorsal Root Ganglia (DRGs) at DIV7 depending on the number of uses of the SBC chip. DRG 𝑛 = 3 − 6, embryos 𝑒 = 3 − 6. The results are presented as the mean±SEM. A Tukey’s multiple comparison test was conducted. The alpha value threshold was set at 5 %, and the *P*-values are represented as follows: ∗ 𝑝 < 0.05, ∗∗ 𝑝 < 0.01, ∗∗∗ 𝑝 < 0.001. **D.** Optical profilometry of the microchannels of a new SBC chip and a SBC chip used four times.

### 2.5. The reversible bonding of the SBC provides direct access to the cells

The reversible bonding properties and detachment of the SBC microfluidic chips also allowed direct access to the cells (**Movie SI.2**). Immunostaining of the cells was performed after the detachment of the device without damaging the cells **(Figure 5A)**. We can clearly observe axons and nuclei of Schwann cells and fibroblasts that had grown and migrated inside the microchannels. NFH is an anti-neurofilaments-heavy chain while SMI312 is staining the neurofilaments-medium chains (NFM) and NucRed the nuclei. Similar images were obtained on fixed cells by SEM after opening of the chips **(Figure 5B)**. The cells were not damaged by the detachment or the drying process required prior to metallization. The reversible bonding properties also allowed direct access to the cells, which could then be scraped in order to realize WB and RT-qPCR (see **Figure 5C-D**). WB was performed after 11 days of culture by scraping DRGs either from a 4-well plate or from a 6-well plate after detaching the SBC chips. Four DRGs were pooled together, and the results showed no significant difference in the expression of SOX10, a protein expressed in Schwann cells, between the DRGs extracted from the control and the SBC devices (**Figure 5C**). RT-qPCR was conducted on a single DRG explant scraped from a well or a chip after 7 days of culture. The comparison of the amount of RNA recovered in control wells and that recovered in the reversibly bonded chips revealed that the SBC chips provide more material under equivalent culture conditions – see **Figure SI.14**. The expression levels of *Krox20*, *Mbp* and *Dhh*, which are markers of the Schwann cell lineage, were found to be comparable in the control and SBC chips (**Figure 5D**). Dhh and Mbp are expressed at the precursor level of Schwann whereas *Krox20* specifically confirms the transition to immature Schwan Cells (prior to myelinating Schwann Cells).^36^

**Figure 5:**
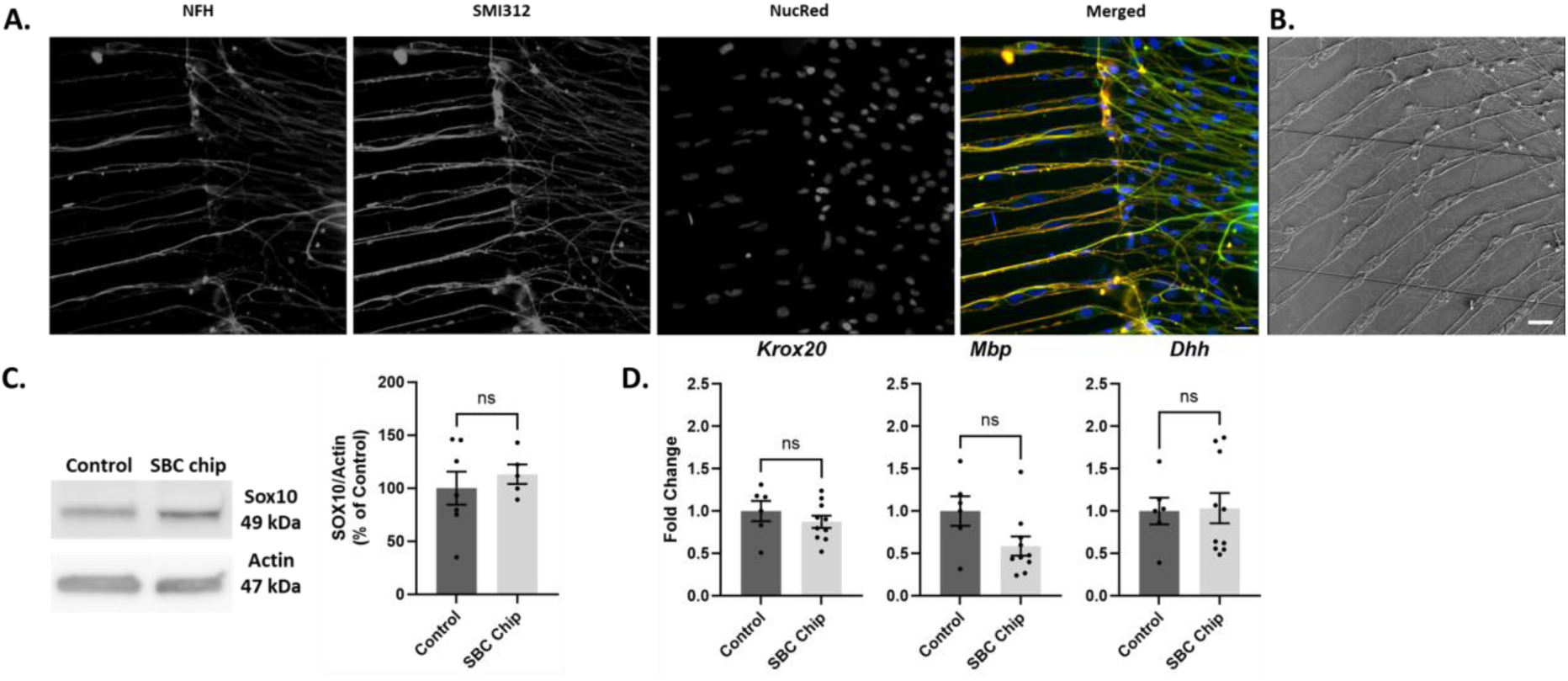
Direct accessibility to cells. **A.** Immunostaining of the neurofilaments of a Dorsal Root Ganglia (DRG) at *DIV*7 after opening of the chip. Nuclei (blue), neurofilaments H (green), SMI312 (red). **B.** SEM image of the neurofilaments of a DRG at *DIV*21. Scale bars represent 20 µ*m* **C.** Western blot analysis of 4 explants DRGs at DIV11 in control wells from a 96-well plate and in Styrenic Block Copolymer (SBC) opened chips and corresponding graph representing Sox10 expression. Actin was used as a loading control. Results represent the mean ± SEM of three independent experiments (𝑛 = 5 − 8, embryo 𝑒 = 3). A t-test was performed for statistical analysis (ns=non-significant). **D.** RT-qPCR analysis of *Krox20*, *Mbp* and *Dhh* genes of one DRG at *DIV*7 in controls and in SBC chips. Results represent the mean ± SEM of three independent experiments (𝑛 = 6 − 10, embryo 𝑒 ≥ 5). A Mann-Whitney-test was performed for statistical analysis (ns=non-significant).

Many reversible bonding strategies have recently been reported, for applications beyond cell culture.^30^ Most of them require the intervention of an external help, a change in the design or the addition of an interlayer between the chip and the substrate. PDMS can be reversibly bonded in the absence of plasma treatment, but it is not easily reproducible and leakage occurs rapidly. In this case, axons can grow under the microchannels rather than inside them and the hydrophobicity of the PDMS, which is usually made hydrophilic by plasma bonding, prevents liquid from easily entering the device.^29^ The self-adhesive properties of thermoplastic elastomers have also been used for reversible bonding^40^ and their reported maximum working pressure without plasma bonding is 3.4 times higher than that of PDMS one.^41^ Recent studies have shown that PDMS can exhibit reversible bonding properties after plasma activation of the substrate, 0,1% APTES treatment and overnight bonding at 75°C^42^. However, the SBC we use has self-adhesive properties that do not require any chemical treatments or design changes. We have shown that it can be reused without impacting the cell culture and no leakage was observed. Axons grew inside the microchannels and the SBC chips were hydrophilized by using plasma treatment without impacting their reversible bonding. Cleaning the chips after use has been suggested before,^43^ but to our knowledge, has never been described. We proposed a fast and simple detailed protocol for cleaning the devices after use, thus enabling them to be reused. However, the reusability of the chips is limited in this particular context. After 3 uses, the microchannels were repeatedly clogged, completely blocking the flow between compartments. We hypothesize that this could be due to the uniform repetition of the different thermal cycles and plasma treatments, as the microchannels clogging was identical and regularly distributed. For chips with thicker channels (in this case the clogged channels were 2.8 µm thick), it is possible to exceed the n = 3 limit.

Axons are not only growing inside the central chamber, but also in the reservoirs (**Figure SI.12**). When the chip is detached, these axons are torn at the same time. However, most of the axons present in the microchannels remain attached. The samples can be immunostained after fixation. The resulting images were more contrasted, with a more uniform background than with the devices, due to the contribution of the autofluorescence of the polymer chip. This direct access also allowed to achieve direct collection for biological analysis of proteins (western blot), which has already been performed using a detachable substrate device,^14^ or nucleic acids (RT-qPCR), which has been carried out directly on the chip.^44^ The detachment of the chips allowed easy scraping of the cells to collect most of the biological material. WB was performed on 4 DRGs pooled, experiments have been conducted on 1, 4 and 8 DRGs in controls to assess the number of DRGs necessary to extract sufficient proteins (**Figure SI.13**). The results showed that there was no significant difference by using 4 or 8 DRGs. However, there is almost 40% of signal intensity difference between the 1 DRG and 4 DRGs experiments, supporting our choice to pull 4 explants together. RT-qPCR was performed by pulling 1 or 2 DRGs explants and similar results were obtained, confirming that enough RNA was collected by scrapping 1 DRG to perform an experiment. Moreover, around 20 𝑛𝑔/µ*L* of RNA was extracted from 1 DRG which would be sufficient to test approximatively 30 genes.

### 2.6. Comparison of carbon footprint of SBC and PDMS chips fabrication process

The energy and time needed to assemble an SBC and a PDMS device were calculated at each stage of the fabrication process **(Figure 6A-B)**. The total energy required to fabricate an SBC chip was 289.3 𝑘𝐽, almost 2 times less than the molding step of a PDMS chip alone, which required 556.8 𝑘𝐽. PDMS chip fabrication took more time and consumed more supplies, resulting in more waste than an

**Figure 6:**
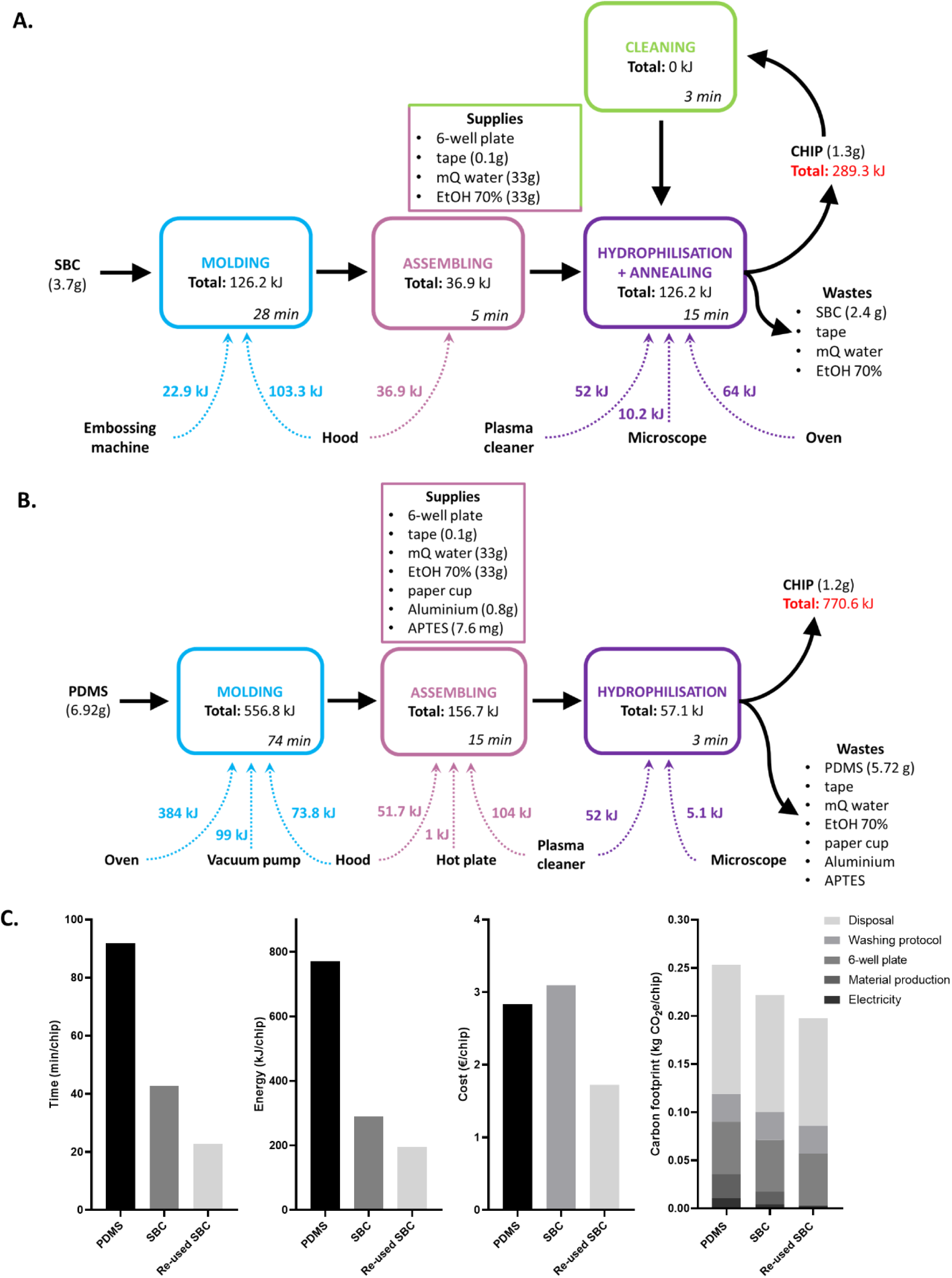
Assessment of the resources used and CO2 emissions generated to produce a Styrenic Block Copoly-mer (SBC) chip and. **a** PolyDiMethylSiloxane **(PDMS) chip.** Energy consumption of the different steps of a fabrication process for **A.** a SBC-based chip and **B.** PDMS-based chip. **C.** Comparison of the time, energy, cost and carbon footprint associated to the fabrication of a PDMS chip, a SBC chip and a reused SBC chip. Note that we did not take into account the labor cost and the equipment maintenance, as these are similar in our context for both materials.

SBC chip. Moreover, an SBC device could be cleaned and reused 2 times before disposal. The cleaning step required no energy and took only 3 min, thereby reducing the time and energy needed for the fabrication process, as well as the cost and kg of CO2 generated **(Figure 6C** and **Table SI.1**). Compared to a PDMS chip, a reused SBC device took 4 times less energy and time to manufacture. The overall price for a single PDMS chip is €2.83, while the price of an SBC chip is €3.10. However, the price of the SBC device decreased to €1.72 when it was reused. To determine the carbon footprint generated by the SBC raw material production, we assumed that it was the average of the carbon footprint of polystyrene, polybutadiene and polyethylene production. The resulting value for one SBC chip is 0.013 𝑘𝑔 CO_2_e while for a PDMS chip it is 0.025 𝑘𝑔 CO_2_e. When an SBC device is reused, as there is no need for new SBC there is no more carbon footprint generated by the material fabrication. The overall carbon footprint has been calculated by considering the mass of CO_2_ equivalent generated per chip for electricity consumption, polymer for chip and 6-well plate production, washing of the chip and disposal of the wastes by incineration. Most of the carbon footprint was attributed to the disposal and washing protocols and was similar for the three types of chips. However, the reused SBC devices produced 0.05 𝑘𝑔 CO_2_e less than the PDMS ones because they consumed less electricity and didn’t require the production of polymer.

Time, energy and carbon footprint generated for SBC and re-used SBC chips fabrication were lower than for PDMS ones fabrication. PDMS required more material and thus generated more waste and cost. Thermoplastics were cheaper to produce^45^ but because PDMS is produced in larger quantities, its final price is lower. It should be noted that the calculations are based on low carbon electricity, coming mostly from nuclear power plants. Depending on the country of reference, electricity can affect both the price and the carbon footprint. Costs and carbon footprint could be lowered for both SBC and PDMS devices fabrication by reducing the amount of waste generated as they cause half of the mass of CO_2_e produced. The quantities of ethanol and milliQ water could be minimized, and increasing the density of chips per mold would also reduce the amount of energy and time necessary to produce them. However, the amount of wasted SBC is easier to reduce than for PDMS because the SBC can be cut to the desired size whereas PDMS must be poured over the entire surface of the mold. When comparing all the environmental impacts from cradle to grave, normalized to those of PDMS, SBC and reused SBC chips appeared to be lower for all impacts, in particular for marine eutrophication and mineral resource scarcity (**Table SI.2**), confirming the interest of using SBC as an alternative to PDMS. In our work, we choose to focus only on the manufacturing process for comparison purposes, we did not take into account the labor cost and the equipment maintenance as they are similar, in our case, for both materials. However, the total energy required to produce one chip can be considered negligible when compared to the energy consumed in the lab or the energy required to produce and transport the material.^46–48^ If we considered the fabrication of 300 chips over a year, the carbon footprint associated with chip fabrication corresponded to 1/60^th^ of the mean carbon footprint of one researcher in one year (6t_eq CO2_).^46^ In summary, the SBC already shows a very interesting reduction in time and cost for the prototyping phase and could have a significant impact on the environmental impact of a product on an industrial scale. This analysis underlines the importance of quantifying and optimizing the environmental impact of microfluidic devices, especially in a field where there is an ongoing industrial scale-up that should intensify with the growing rationalization of animal models.

## 3. Conclusion

We introduced a detachable, reusable, easy to produce and equivalent in cell culture performance SBC device for an original application to DRGs culture in the context of peripheral nervous system study. We proposed a new design adapted to DRGs explants and their radial growth with 6 peripheral compartments. We estimated the shear stress in the chip using simulations and particle velicometry and experimentally confirmed its ability to perform axonal growth and viability assays of a murine peripheral nervous system model. Based on the above biological assays, the PDMS devices, used as a control, showed equivalent performance to the SBC devices. The demonstration of growth and maintenance of a primary co-culture of nerves, which are particularly sensitive to stress, in an SBC chip paves the way for tissue diversification and species growth in SBCs for the neurofluidic community, but more generally for the microphysiological systems community. The reversible bonding of the chips has been exploited both for direct access to different contact imaging microscopy and biological analysis techniques such as SEM, WB and RT-qPCR. The possibility to recycle our chips led us to develop an original workflow to evaluate the resources (time, energy, financial costs and CO2) involved in the production and recycling of the chip, which could be used by the microfluidic community in general to initiate a deeper rethinking. Switching to a reusable SBC chip allowed a significant reduction in all these resources, more than 4 times in time and energy compared to PDMS, contributing to more sustainable research. The carbon footprint assessment methodology can be reused for other devices. In the field of the microphysiological systems, the paradigm to move away from PDMS toward soft thermoplastics makes it possible to reduce the amount of resources used and to overcome the usual limitations for cell culture. This could help minimize the environmental impact of a rapidly growing industry. It also contributes directly to efforts in microfluidics to accelerate the clinical translation of therapeutics for neuropathologies.^49,50^ From an animal ethics perspective, the development of alternative *in vitro* models has the potential to significantly reduce the animal use, in alignment with the 4Rs commitment. In the case of DRGs, the number of embryos in a pregnant mouse is typically between five and seven, with an average of 40 DRGs per embryo. This yields a potential 250 experiments to be conducted from a single mouse, depositing one dorsal root ganglion per chip. This approach allows for the testing of up to 1,500 conditions using the proposed design, rendering it highly valuable for drug screening and mechanistic investigation studies. Finally, the demonstration of viable DRG culture on our chip paves the way toward myelinating coculture in the PNS, which would provide 2D myelin models for neuropharmacological explorations. This could be used to study of the processes of myelination, demyelination, and remyelination, but also to select *in vitro* drugs that could be used for regenerative treatments by stimulating the production of new myelin.

## 4. Experimental Section

### 4.1. Chip design and fabrication

#### a) Molds fabrication

The photomask designs were drawn using CleWin 5.2.4 version and printed on a SodaLime substrate with a resolution of 128k dpi (JD Photo Data, UK). A first layer of 2.8 µm thick microchannels was structured with SU-8 2002 (Chimietech, France) on a 4-inch diameter silicon wafer 1 *mm* thick using soft lithography. This was followed by the addition of a second layer of 100 µ*m* thick chambers made with SU-8 2050 (Chimietech, France). The Epoxym Kit (EdenTech, France) was used to create a negative replica in 1: 10 Sylgard 184 PDMS (Neyco, France), followed by an epoxy positive replica of the SU-8 mold with Gallon KIT Conapoxy FR-1080 (Ellsworth Adhesives, USA).

#### b) Thermomolding and assembly of SBC chips

The microfluidic chips were embossed from styrene block copolymer (SBC) foils (EdenTech, France) at 160 °𝐶 for 3 *m*𝑖𝑛 (0.07 − 0.08 *MPa*) using the Sublym hot press machine (EdenTech, France) and the previously fabricated epoxy mold. The central chamber was punched with a 3 *mm* punch to obtain an open top device. A second layer of SBC embossed on a flat epoxy mold was added to each chip with a 6 *mm* diameter reservoir for the central chamber. The outer chambers were punched simultaneously in the patterned layer and the reservoir layer using a 4 *mm* puncher. Finally, each chip was die-cut with a 26 *mm* puncher. The two-layer devices were then placed in a 6-well plate (VWR, USA) for hydrophilization by plasma treatment for 10 *m*𝑖𝑛 in a plasma cleaner/sterilizer (Harrick Scientific Products, USA). They were baked at 80 °𝐶 for 2 ℎ to increase the bonding strength.

#### c) Fabrication and assembly of PDMS chips

Polydimethylsiloxane (PDMS) chips and reservoirs were respectively molded with 1:10 Sylgard 184 from the SU-8 mold and a basic silicon wafer, and baked at 80 °𝐶 for 2 ℎ𝑜𝑢*rs*. The patterned layer and the reservoir layer were punched with the same punchers used for the SBC chips and both were plasma treated for 10 *m*𝑖𝑛 prior to assembly. 6-well plates were pre-treated with 1 % (3-aminopropyl)triethoxysilane (APTES, Sigma-Aldrich, Germany) in milliQ water. Finally, the devices were treated with plasma again for 10 *m*𝑖𝑛 and were bonded in the 6-well plates.

#### d) Sterilization and coating of the chip

The same protocol was used on the SBC and PDMS chips. The plates containing the devices were placed in a UV/Ozone ProCleaner™ Plus (BioForce Nanosciences, USA) for 10 *m*𝑖𝑛 for sterilization. The chips were then rinsed with EtOH 70 % and three times with Dulbecco’s phosphate buffered saline 1X (D-PBS 1X, Gibco, USA) before being incubated with 5 % Poly-L-lysine (Sigma-Aldrich, Germany) in a solution of boric acid (Sigma-Aldrich, Germany) and sodium hydroxide for 30 *m*𝑖𝑛. at 37°C. After washes with Neurobasal medium (Gibco, USA), they were coated with a solution of 130 µ𝑔. *m*𝑙 − 1 rat tail collagen I (Clinisciences, France) in 17.5*mM* acetic acid for 1 h at room temperature. Finally, the chips were rinsed again and placed at 37 °𝐶 and 10 % CO_2_ overnight before cell seeding. The same coating procedure was followed for the 4-well and 96-well plates, which were used as controls.

#### e) Detachment and cleaning of the chip

The wells were filled with 1 *mL* of D-PBS to immerge the chips before to progressively detach them with a tweezer. After 10 *m*𝑖𝑛 of incubation at room temperature, the chips were carefully removed from the wells. If cells observation was not needed, the devices were detached directly without the incubation step. To clean them, they were first bathed in a solution of milliQ water and soap solution (disinfectant Dermanios Scrub CG, Laboratoire Anios, France) and gently brushed with a toothbrush, then rinsed successively with milliQ water, EtOH 70 % and with milliQ water again. Finally, they were either dried in a laminar flow hood for 1 ℎ before storage or dried with compressed air and directly reused following the usual protocol of sterilization and coating. Optical profilometry was performed with a WykoNT9100 optical profilometer (Veeco, USA) on a new chip and a chip that had been used up to 4 times and cleaned each time.

### 4.2 Dorsal Root Ganglia (DRG) Cell Culture

#### a) Cells harvesting, seeding and maintenance

Dorsal root ganglia (DRGs) were harvested from embryos at *E*13.5 surgically removed from pregnant C57BL/6 J mice (Janvier, France) following a previously described protocol.^51^ For explant culture, the devices were filled with DRG plating media (DMEM high glucose (Gibco, USA), Glutamax 1X (Gibco, USA), 10% Heat Inactivated Horse Serum (Gibco, USA), 2.5S NGF (R&D Systems, USA), Antibiotic/Anti-mycotic 1X (Gibco, USA)). One DRG was placed in the central chamber of each microfluidic chip or in the center of a well of a 4-well plate. For dissociated cell culture, the DRGs were trypsinized with a solution of 0.25 Trypsin in Hanks’ Balanced Salt Solution 1X (HBSS 1X, Gibco, USA) for = at 37 °𝐶 and 5 % CO_2_. Trypsinization was stopped with L-15 media (Gibco, USA) containing 10% horse serum. DRGs were then spun down at 1,500 *r*𝑝*m* for 5 *m*𝑖𝑛. The cells were resuspended in DRG plating medium and seeded at a targeted density of 275 000 *c*𝑒𝑙𝑙*s*. *cm*^−2^ inside the central chambers or in a well from a 96-well plate, as a control. The dissociated cells were seeded in a volume of 7 µ*L* in the central chamber and incubated 2 ℎ prior to fill the reservoir, to allow them to adhere to the surface of the culture plate. DRGs were incubated at 37 °𝐶 and 10 % CO_2_. 18 to 20 ℎ after the seeding, the medium was replaced with Neurobasal medium supplemented with Glutamax 1X, 2.5S NGF, B27 Supplement (Gibco, USA) and Antibiotic/Antimycotic 1X. The medium was changed every 2 𝑑*a*𝑦*s*. To prevent evaporation, 3 *mL* of D-PBS was added in between the wells of the plates. All aspects of animal care and animal experimentation were performed in accordance with the relevant guidelines and regulations of INSERM and Université Paris Cité (authorization APAFIS#7405-2016092216181520).

#### b) Live and dead cell assays

Viability was assessed at day *in vitro DIV*3 and *DIV*7 using a LIVE/DEAD Viability/Cytotoxicity Assay Kit (Invitrogen, USA). Cells were incubated for 30 *m*𝑖𝑛 at 37 °𝐶 and 10 % CO_2_ and rinsed with L15 medium without red phenol (Gibco, USA). Cells were observed using the Cy5 and EGFP channels on a ZEISS Axio Observer Z1 microscope (Zeiss, Germany) incubated at 37 °𝐶 and 10 % CO_2_ and the images were processed using ImageJ 1.54j.^52^ The data obtained were analyzed using R and tidyverse and the graphs were plotted on GraphPad Prism 8 (GraphPad Software, USA). The full R markdown script is available as supplementary information (*Script SI.1*). Sidak’s multiple comparison tests were performed. The alpha value threshold was set at 5 % and the *P*-values are presented as follows: ∗ *P* < 0.05, ∗∗ *P* < 0.01, ∗∗∗ *P* < 0.001.

#### c) Immunostaining

Cell cultures at *DIV*3 and *DIV*7 were fixed with 4% paraformaldehyde (PFA, Electron Microscopy Sciences, USA) for 10 *m*𝑖𝑛 at room temperature. At *DIV*7, the SBC chips were detached to facilitate the staining. The cells were then permeabilized (0.25 % Triton X100 (Sigma-Aldrich, Germany) and 0.1 % Tween20 (Thermo Fischer Scientific, USA) in D-PBS) for 10 *m*𝑖𝑛. After washes with D-PBS and incubation for 45 *m*𝑖𝑛 in a blocking solution (0.5 % Triton X100, 0.1 % Tween20, 2 % bovine serum albumin (BSA, Sigma-Aldrich, Germany), 3 % Mouse on Mouse blocking reagent (Bio-Techne, USA) and 5 % Normal Donkey Serum (Sigma-Aldrich, Germany), the cells were incubated overnight at 4 °𝐶 with the primary antibodies: rabbit anti-NFH (1: 1000, Merck Millipore Ref. AB1989, Germany) and mouse anti-SMI312 (1: 250, BioLegend Ref. AB_2566782, USA). The cultures were again rinsed with D-PBS and incubated for 1h at room temperature with the secondary antibodies: donkey anti-rabbit Alexa Fluor™ 488 (1: 1000, Invitrogen Ref. A-21206, USA) and donkey anti-mouse Cyanine3 (1: 165, Jackson Ref.

AB_2340813, USA). For nuclear staining, the devices were incubated 30 *m*𝑖𝑛 in a solution of NucRed (Invitrogen, USA) in HBSS. To reduce the amount of antibody and staining required for chip detachment, rings were 3D printed in polylactic acid and placed in each well to reduce the surface area (**Figure SI.1-2**). Images were captured on a ZEISS Axio Observer Z1 microscope and processed on ImageJ using the *NeuriteJ* plug-in. The resulting data were processed in RStudio (Posit, USA) and graphs plotted on GraphPad Prism 8. The full R markdown script is available as supplementary information (**Script SI.2**) Sidak’s multiple comparison tests were performed.

#### d) Western Blot

Western blots (WB) were performed on DRGs explant cultures at *DIV*11. The chips were detached, and the proteins were extracted by scraping the DRGs in RIPA buffer (50 *mM* Tris Base, 150 *mM* NaCl, 1 *mM* MgCl_2_, 1 % NP-40 (Sigma-Aldrich, Germany), 0.1 % sodium dodecyl sulfate SDS (EU0660, Euromedex, France), 0.5 % sodium deoxycholate (D6750, Sigma-Aldrich, Germany), Protease Inhibitor Complete mini EDTA free (11836170001, Roche, Switzerland), Pyrophosphate (10 *mM*, S9515, Sigma-Aldrich, Germany), Phosphate Inhibitor Cocktail 2 (10 µ*L*/*m*𝑙, P5726, Sigma-Aldrich, Germany), Phosphate Inhibitor Cocktail 3 (10 µ*L*/*m*𝑙, P0044, Sigma-Aldrich, Germany) and sodium fluoride (10 *mM*, S7920, Sigma-Aldrich, Germany). The protein quantification was determined using a Bio-Rad DC Protein Assay (Bio-Rad, USA) with BSA as the standard. A total of 15 µ𝑔 of protein was loaded into a 4 − 20 % TGX 10-wells gel (Bio-Rad, USA) and the transfer step was performed using a Trans-Blot Turbo transfer system (Bio-Rad, USA) on polyvinylidene difluoride membranes (Bio-Rad, USA). Non-specific binding sites were blocked with blocking solution (5 % BSA in Tris Buffered Saline (TBS, Euromedex, France)) and then incubated with primary antibodies rabbit anti-Sox10 (1: 1000, ab155279, Abcam, UK) and rabbit anti-Actin (1: 2 000, ab8227, Abcam, UK), followed by a goat anti-rabbit Horseradish Peroxidase (HRP) coupled secondary antibody (1: 20 000, 111-035-144, Jackson IR, UK). Finally, specific bands were detected with the Pierce®ECL 2 Western Blotting substrate kit (RPN2209, Thermo Fischer Scientific, USA), using the ImageQuant LAS 4000 ECL imaging system (GE Healthcare, USA). The images were analyzed using the gel tool analyzer of ImageJ (1.54g). The integrated densities of Sox10 were normalized to β-actin expression and plotted using GraphPad Prism 10. A t-test was performed for statistical analysis.

#### e) RT-qPCR

RT-qPCR was performed on explants at *DIV*7. The chips were detached and the DRGs were scraped in Trizol (15596018, Invitrogen, USA). The RNA was precipitated and resuspended in water prior to quantification with a Nanodrop (Thermo Fischer Scientific, USA). The SuperScript III Reverse Transcriptase (18080085, Invitrogen, USA) was used following the manufacturer’s instructions to realize the reverse transcriptase of 90 𝑛𝑔 of RNA. qPCR was performed on a CFX384 biorad qPCR machine (15 *s* of denaturation at 95 °𝐶, 30 *s* of elongation at 60 °𝐶 – 40 times) by using Sybr Green Rox kit (AB-1162/B, Thermo Fischer Scientific, USA) containing 300 𝑛*M* of selected primers (Table 1). The quality of the qPCR was tested by checking the dissociation curve. Each qPCR reaction was performed in triplicates. Relative expression of target genes was normalized to Ppia and Rpl13a mRNA levels and calculated using the ΔΔCt. Results are expressed as fold changes with respect to the controls and the data were plotted using GraphPad Prism 10.

**Table 1.**
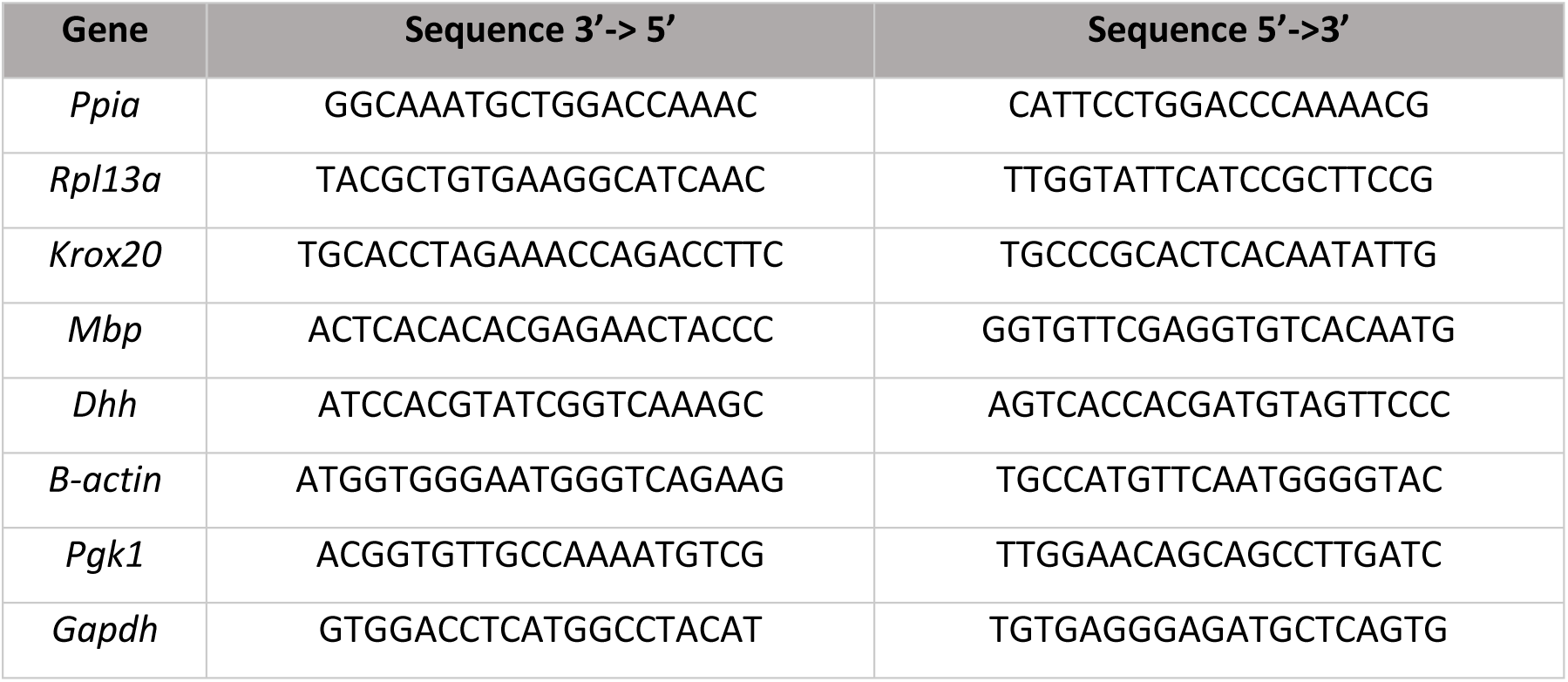
qPCR primers used in this study.

#### f) Scanning Electron Microscopy

After 21 𝑑*a*𝑦*s* of culture, the cells were rinsed with D-PBS and fixed in 4 % PFA. The bottom of each well was then laser-cut using a Universal Laser Systems VLS 4.60 (Universal Laser Systems, Inc., USA) to obtain circular slides suitable for scanning electron microscope. The chips were detached according to the previous protocol and the biological samples were dried using an automated critical point dryer Leica EM CPD300 (Leica Microsystems, France). The samples were then metallized with platinum in a Cressington 208HR metallizer (Cressington Scientific Instruments, UK) and observed in a Zeiss Gemini SEM 360 scanning electron microscope (Zeiss, Germany).

#### g) Finite Element Method (FEM)

COMSOL Multiphysics software was used to perform diffusion-convection simulations on the microfluidic device. The *creeping flow* and *transport of diluted species* modules were implemented for this study. In the diffusion-convection experiment, a pressure gradient of 5 and 20 *Pa* between the inlet and the outlet was used for the convective flow, and an inflow of 0.5 mM of water (diffusion coefficient 𝐷 = 10^−9^ *m*^2^. *s*^−1^) was used for diffusion, by keeping the inlet and outlet the same (transport) or opposite (isolation). The system comprises two identical blocks of size 1 × 1 × 0.1 *mm*^3^, merged with a smaller block of size 0.5 × 1 × 0.028 *mm*^3^. This geometry represents the junction between the central and peripheral chambers, which is of particular relevance for the study. Approximate solutions of the following equations were numerically calculated by the FEM software:

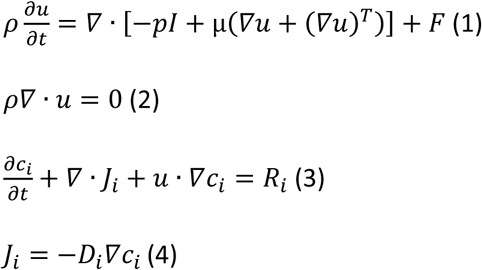

Equation (1) is the Navier-Stokes equation, (2) is the incompressibility of the flow, (3) is the diffusion-convection equation, and equation (4) is Fick’s law of diffusion with 𝜌 the fluid density, 𝑢 the fluid velocity, 𝑝 the fluid pressure, 𝐼 the identity matrix, 𝑇 the temperature, µ the fluid dynamic viscosity, 𝐹 the external forces applied to the fluid, and where, for species 𝑖, *c*_𝑖_ is the concentration, 𝐽_𝑖_ is the molar flux, 𝑅_𝑖_ is a net volumetric source for *c* and 𝐷_𝑖_ is the diffusion coefficient.

#### h) Particle image velocimetry

The devices were first coated with 5% BSA in D-PBS1X for 10 min to reduce collision impact with the walls. FluoSpheres^TM^ amine (0.2 µ*m* diameter) (Invitrogen, USA) were diluted 1/40000 in the same solution. To obtain a flow of 25 *Pa*, 60 µ*L* of the microbead solution was injected into the central chamber and 5 µ*L* of the solvent solution was injected into the peripheral chambers. D-PBS was added in between the wells to prevent evaporation and the plate was sealed with parafilm. Microbeads were observed using the Cy5 channel on a ZEISS Axio Observer Z1 microscope. Images were analyzed directly using the Zeiss Zen software and data were plotted using GraphPad Prism 8.

#### i) Time, cost, carbon footprint and energy consumption calculations

The estimated times, costs, carbon footprint and energy consumption were calculated for one chip. Power consumption was measured using a wattmeter Primera-Line PM 231 E (Lextronic, France). The measurements were converted to energy by multiplying the power by the time of use of each piece of equipment. Whenever an equipment was used for several devices at the same time, the amount of energy and time were divided by the number of chips to estimate the amount required for one device. Material prices were based on supplier prices. Electricity prices were based on the French electricity price in June 2024 (€0.2516/𝑘𝑊ℎ). The carbon footprint was calculated based on the direct emissions of products manufactured in Europe, using the Ecoinvent 3.5 database, required for the fabrication of one chip. PDMS production was assimilated to silicone using the Ecoinvent process ‘product production | silicone product | APOS, S RER’, while SBC manufacture was modeled as an average of 25 % polyethylene, 25 % polybutadiene and 50 % polystyrene using the processes ‘polyethylene production, high density, granulate | polyethylene, high density, granulate | APOS, S, RER’, ‘polybutadiene production | polybutadiene | APOS, S, RER’ and ‘polystyrene production, general purpose | polystyrene, general purpose | APOS, S, RER’. In addition, the impact calculations included electricity consumption, 6-well plate production, washing protocol and waste disposal by incineration. Impacts were calculated using the ReCiPe 2016 Midpoint (H) method.

## Data availability

The data for this article plots can be found on https://github.com/HugoSalmon/25Moreau.

## Supporting Information

Supporting Information is available from the Wiley Online Library.

## Author Contributions

H.S., S.B., S.M., A.S. and A.E-T conceptualized the study. H.S., A.S. and C.D-B. provided the resources. S.M., H.S, A.S. and A.E-T made data curation. S.M., A.S. and A.E-T realized the formal analysis. H.S. and S.B. supervised the study. H.S. acquired the fundings. H.S., T.EJ., A.S. and S.B. validated the study. Investigation was conducted by H.S., S.M., R.F-B., A.S. and C.D-B.. S.M. wrote the original draft and H.S., T.EJ., A.S., C.D-B. and S.B. review and edited the paper. H.S., S.B., S.M., T.EJ., R.F-B., A.S. and C.D-B. designed the methodology of the study. H.S. and S.B. handled the project administration.

## Acknowledgements

S.B. and H.S. contributed equally to this work. This research was funded by the French National Research Agency (ANR-23-CE19-0007-01) and the Université Paris Cité via the Initiative d’Excellence Idex Emergence (IDXD7-MOCHI). The motorized microscope and the motorized macroscope were funded by DIM BIOCONVS Région Ile de France (HCS-Mochi and MisoSoup projects, respectively). RFB was funded by DIM ELICIT Région Ile de France (FOMO project). We thank L. Réa from UMR 7057 MSC for prototyping the microscope adaptor.

The parts were fabricated at the mechanical workshop core facility of BioMedTech Facilities Université Paris Cité (https://biomedicale.u-paris.fr/biomedtech-facilities/) INSERM US36 | CNRS UAR2009 by T. Bastien and V. Pierrat.. The authors thank the staff of UMR 7162 MPQ clean room, the IPGG platform from PSL university for the easy access to the platform and ITODYS platform from Université Paris Cité for the SEM images. Finally, the authors thank V. Sundaram and C. Massaad for the discussions regarding the peripheral nerve modeling, J-M Di Meglio and Dimitri Ayollo for discussions on microscopy.

The authors thank the Mathematics and IT department of Université Paris Cité for the access to https://pleiade.mi.parisdescartes.fr/. The large language model “athene-v2” was used for improving the English of the manuscript.

## List of abbreviations

4R: Reduction, Replacement, Refinement and Responsibility
CNS: Central Nervous System
DIV: Days *in vitro*
DRG: Dorsal Root Ganglia
PDMS: PolyDiMethylSiloxane
PNS: Peripheral Nervous System
RT-qPCR: Reverse Transcription quantitative Polymerase Chain Reaction
SBC: Styrenic Block Copolymer
SEM: Scanning Electron Microscope
sTPE: soft Thermoplastic Elastomer
WB: Western Blot

## 1 Code

R markdown code for automated data treatment can be found here.

- Script SI.1: R markdown script for live-dead data treatments.
- Script SI.2: R markdown script for axonal growth data treatments.

The github depot also contains a data set to run the markdown code on example datasets. Script S1.1 requires ’:LiveDead_Noise_NucGreen.csv”, “LiveDead_Noise_NucRed.csv”, “LiveDead_NucGreen.csv” and “LiveDead_NucRed.csv”. Script S1.2 requires “AxGrowth.csv”.

## 2 Profilometry

We controlled our mold and replica using a mechanical profilometer - see Figure SI.1. The SU-8 on silicon primary mold and the epoxy mold are the most rigid and offer the best contrast. They confirm the thickness of the microchannels is 2.70 *−* 2.80*µm*. 1t is in good agreement with the PDMS and SBC replicas which are 2.65*−*2.78*µm* thick. The parabolic baseline of the PDMS profile is most likely due to the curvature of the 4 inch PDMS replica (*S* = 157.0*mm*^2^) which have thicker edges. The SBC does not display deformation because the replicas are smaller (*S* = 12.3*mm*^2^). The apparent roughness of the SBC hills might be due to the adhesion forces of the polymer (SBC is a pressure sensitive adhesive) which might generate instability. This was not observed when doing optical profilometry (Figure 4D).

**Figure SI.1:**
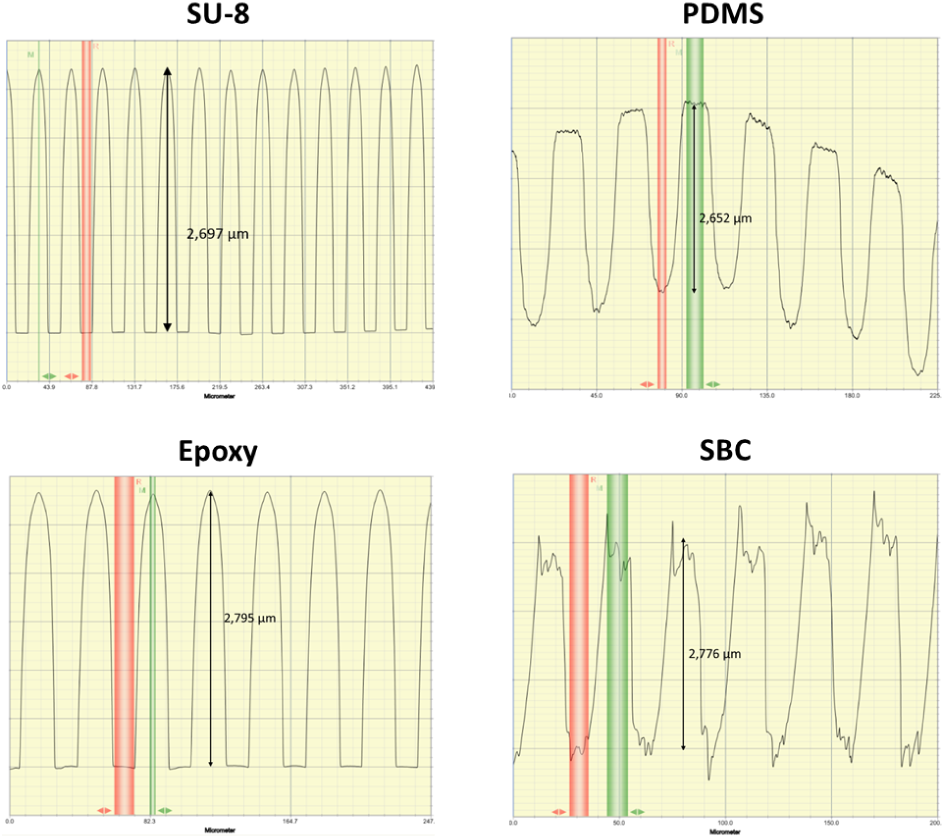
Mechanical profilometry of microchannels of the SU-8, PDMS and epoxy molds and SBC chips. The irregularities observed on the PDMS and SBC profilometers are due to the elasticity of the materials. The height of the microchannels is similar for each replicate.

The punching quality varies greatly between a handpunch on PDMS and a manual cutting press on a flexdym reservoir layer-see Figure SI.2.

**Figure SI.2:**
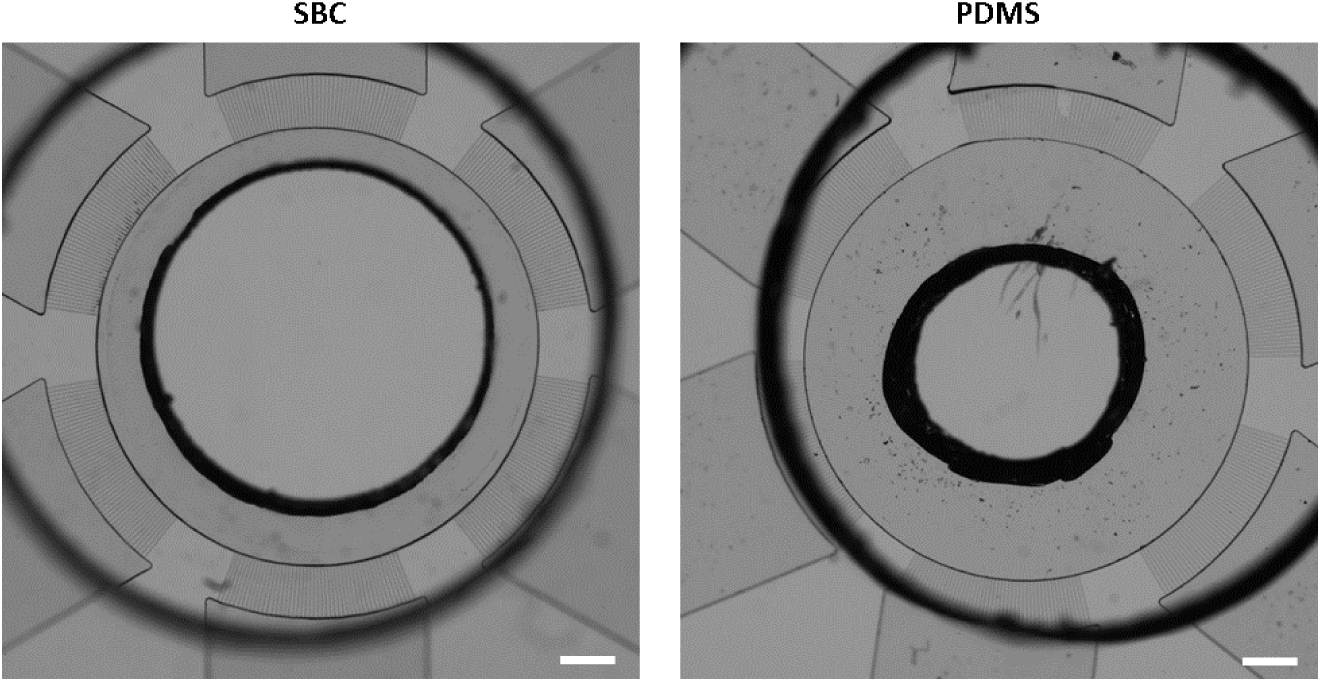
Differences in hole size after punching an SBC and a PDMS chip with a 3 mm puncher. Scale bar represents 500 _µ_m.

## 3 Volume effect on pressure

Evaluating the evolution of the hydrostatic pressure-induced flow Our mathematical model provides an estimation of the flow time inside a microfluidic device. To model the pressure-gradient induced flow between chambers, we simplified the overall geometry of the problem (Figure SI.3.A, 3.B). The hydrodynamic resistance of the equivalent microchannel was calculated using electrical circuit analogy of *n* microchannels of resistance *R_h_* in parallel. Using the conservation of flow rate inside the system, we obtained the following simple differential equation governing the volume of fluid inside a given chamber over time :

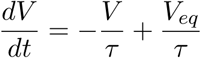

For one cylindrical central chamber of section *S*_0_ and *N* peripheral chambers of same section *S*_1_, the time constant of our model is :

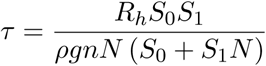

From this formula, we conclude that the time scale depends on the geometry of the device and the density of the fluid, but not on the initial volume conditions. We calculated the numerical solution using water parameters for the fluid, *n* = 27, *N* = 6, *S*_0_ = 2.1 *×* 10*^−^*^5^ m^2^ and *S*_1_ = 1.3 *×* 10*^−^*^5^ m^2^ (Figure SI 3.C, SI 3.D). For these values, *τ ≈* 16h, meaning it takes each chamber about 50 hours to reach 95% of its final volume.

**Figure SI.3:**
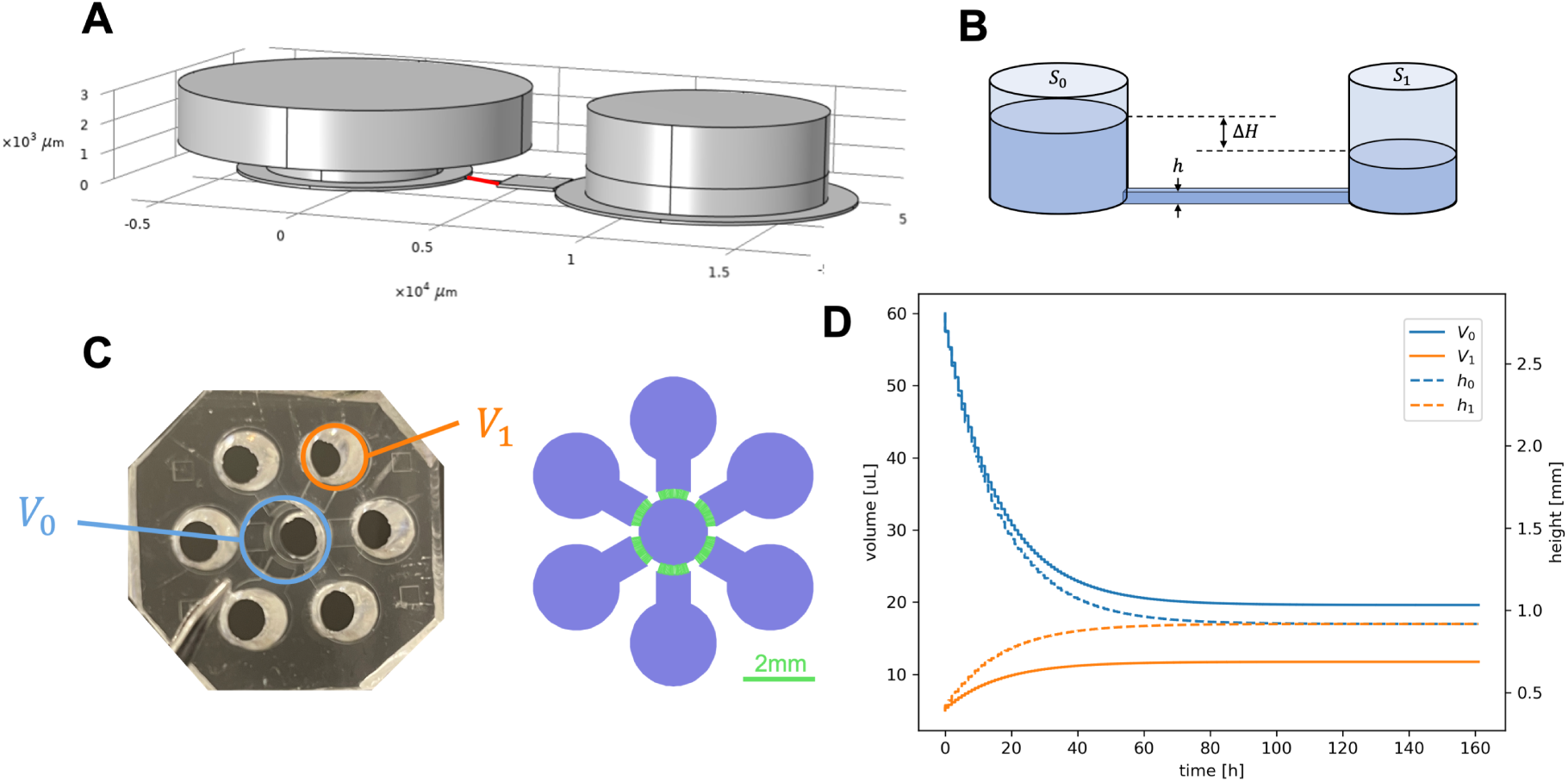
Model of the flow generated inside a microfluidic chip. (A) CAO perspective view of the central chamber (left) and a peripheral chamber (right) connected by one microchannel (in red). (B) Schematic illustration of the flow between two chambers using one-cylinder approximation. (C) Photography of microfluidic chip (left) and schematic illustration of the model used (right). The chambers are shown in purple, and the microchannels in green. (D) Evolution of liquid volume and height inside central chamber and peripheral chamber with respective initial volumes of 60 _µ_L and 5 _µ_L.

The following values were obtained for a pressure gradient of 30Pa, which is the theoretical maximum pressure that can be applied inside the studied microfluidic device given the finite volume of the reservoirs (Figure SI.4). The maximum shear stress inside the cross section is 0.8 dyn/cm^2^. This is lower than the cell viability limit for cortical neurons (5 dyn/cm^2^) which was measured for a 24h exposition time [1J, meaning the fluid flow has no degrading effect on the studied axons.

**Figure SI.4:**
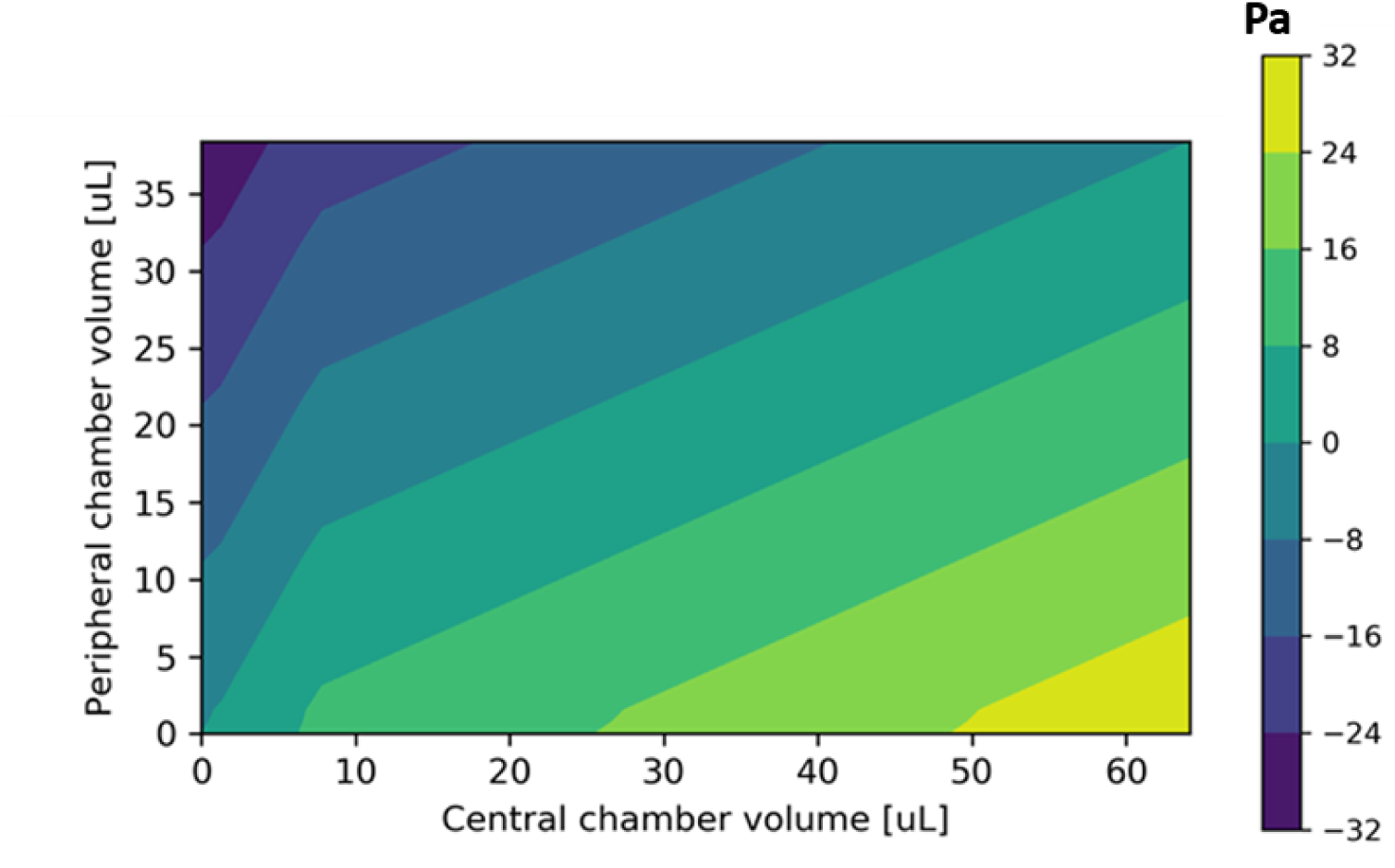
2D graph of the pressure within the chip depending on the volume of fluid in the central chamber and in the peripheral chambers.

Cylindrical approximation According to Pascal’:s law, a pressure-gradient force is applied when two compartments of the same liquid have different altitudes. To obtain the volume as a linear function of the altitude (*V* = *hS*) and thus facilitate calculations, we used a one-cylinder approximation for the chambers. 1n reality, the microfabrication process requires that the chambers be composed of three juxtaposed cylinders. The simplified chambers were calculated by conserving the same height and volume than those of the original ones.

Equivalent circuit theory Microchannels act as hydraulic resistors. The hydrodynamic resistance of a microchannel can be obtained thanks to a well-known approximation for rectangular cross-section channels [2]:

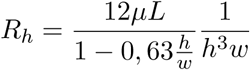

Given the small dimensions *h* = 2.8 _µ_m, *w* = 10 _µ_m and *L* = 500 _µ_m, the obtained value *R_h_* = 3.3 *×* 10^16^ Pa.s.m*^−^*^3^ is very high compared to that of the chambers (about 10^11^ higher). Considering this the only source of flow resistance is therefore a reasonable assumption. Using the conservation of flow rate in the parallel coupling of *n* microchannels, the inverse-additive law gives us the total resistance as 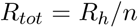 (Figure SI.5). This approximation is true in the case of small Reynolds numbers, which is verified inside our microfluidic device due to the small length and slow speed rate : 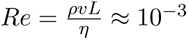.

**Figure SI.5:**
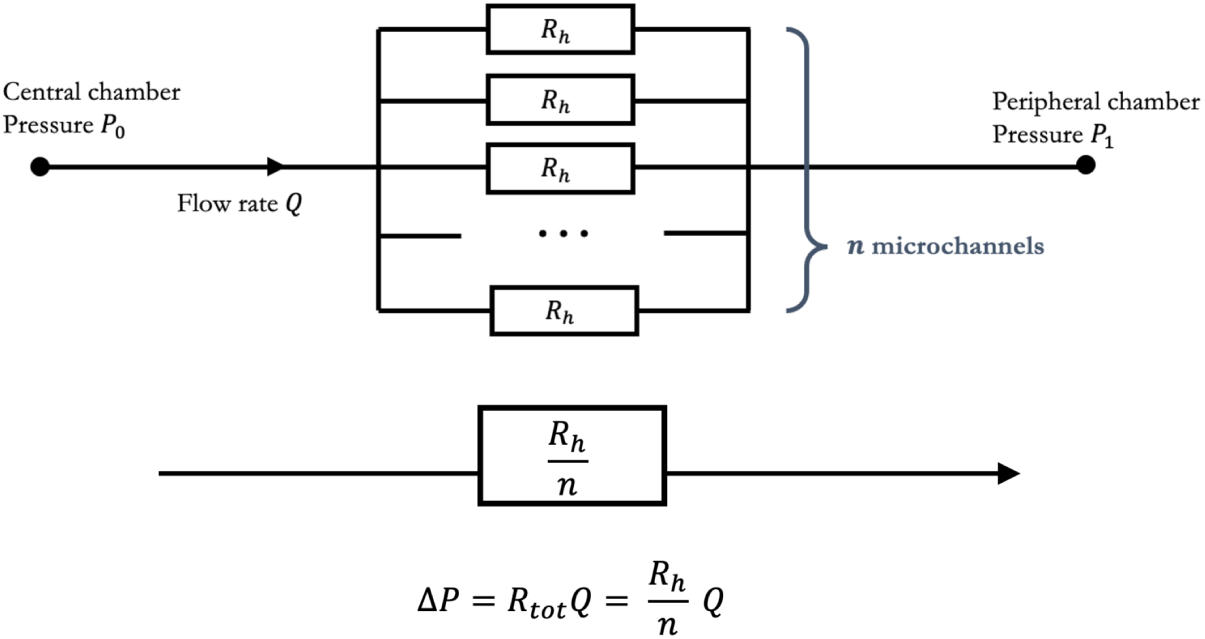
Equivalent circuit theory for two chambers

System equations The definition of the flow rate *Q* between the central chamber (volume *V*_0_) and one of the *N* identical peripheral chambers (volume *V*_1_) can be written as :

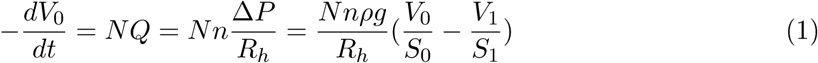

In addition, the total volume of fluid is constant over time :

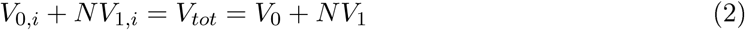

Injecting (2) into (1) gives us the desired differential equation on the volume of either the central or one of the peripheral chambers. This equation was numerically solved using the solve_ivp function from the python library using Runge-Kutta method. The average flow speed inside the microchannels can be deduced since *v* = *Q/hw*.

## 4 Image analysis workflow

The overall study relies on a defined workflow to avoid inconstitency and user dependent analysis. 1t also aims at quantifying exhaustively in a high content manner the overall information available on the chip regarding viability Figure SI.6 and axonal growth Figure SI.7.

As observed in the Figure SI.9, the flower design displays a gradient in each arm that is visible when doing an immunostaining. Any treatment to the axons displays this gradient distribution.

The recycling process after several use demonstrate occlusion next to the central part specifically. Though we did not perform any composition analysis of these occlusion, one hypothesis is that the thermal history of the polymer (long term incubation, cleaning and reconditioning 3 times) lead to a deformation of the smallest patterns.

**Figure SI.6:**
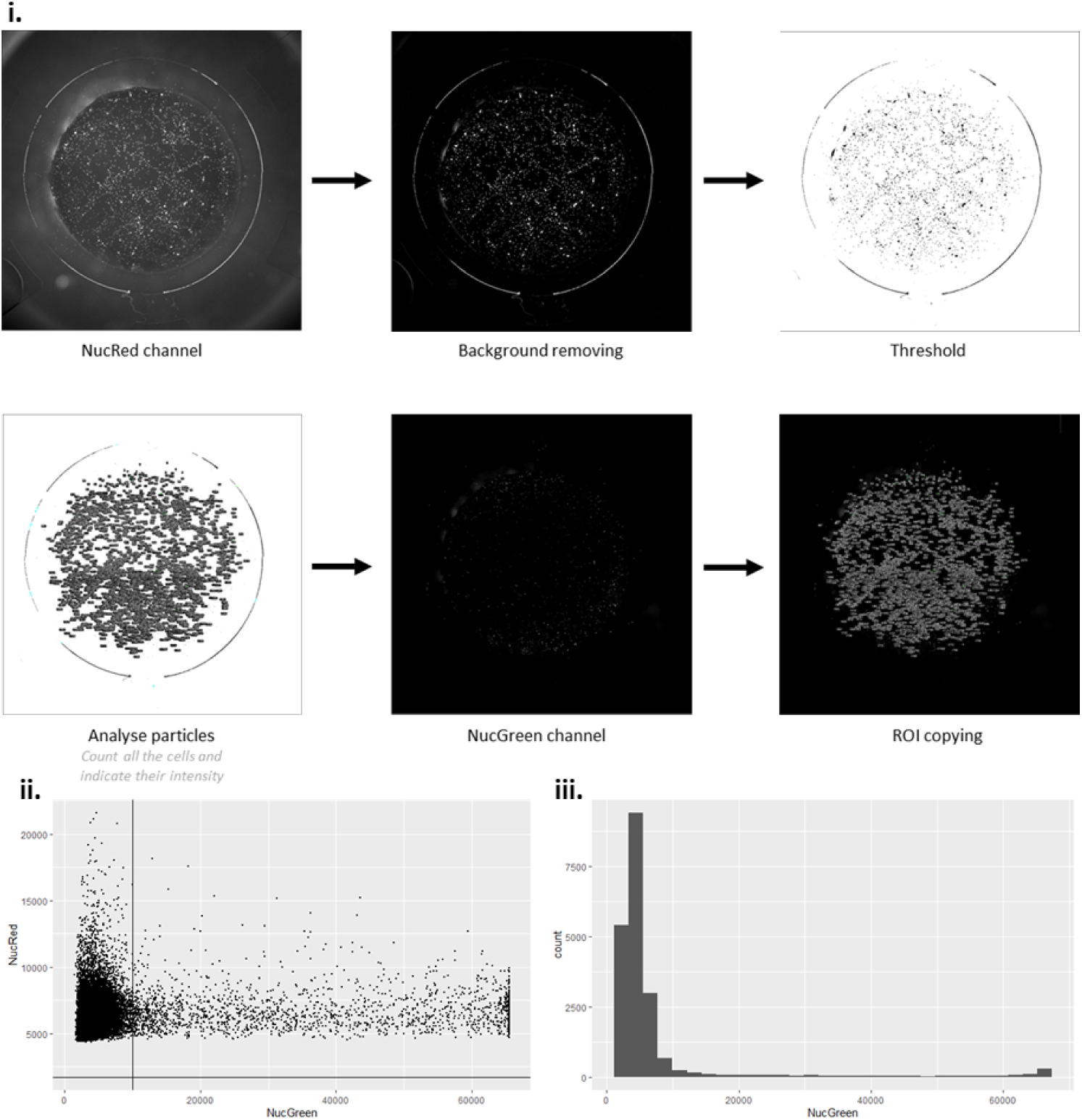
1mage treatment for total live-dead analysis. i. Workflow: The entire chip is scanned as a single tile. The background is removed and the particles are first threshold and counted on the NucRed channel. The white intensity is then compared between the NucGreen and the NucRed channels. From this data, a dotplot ii. and a histogram iii. are plotted with the intensity of NucRed compared to the intensity of NucGreen. The number of living and dead cells is calculated based on the results with a count of living cells ranging from 0 to 10000*UI* for NucGreen intensity.

**Figure SI.7:**
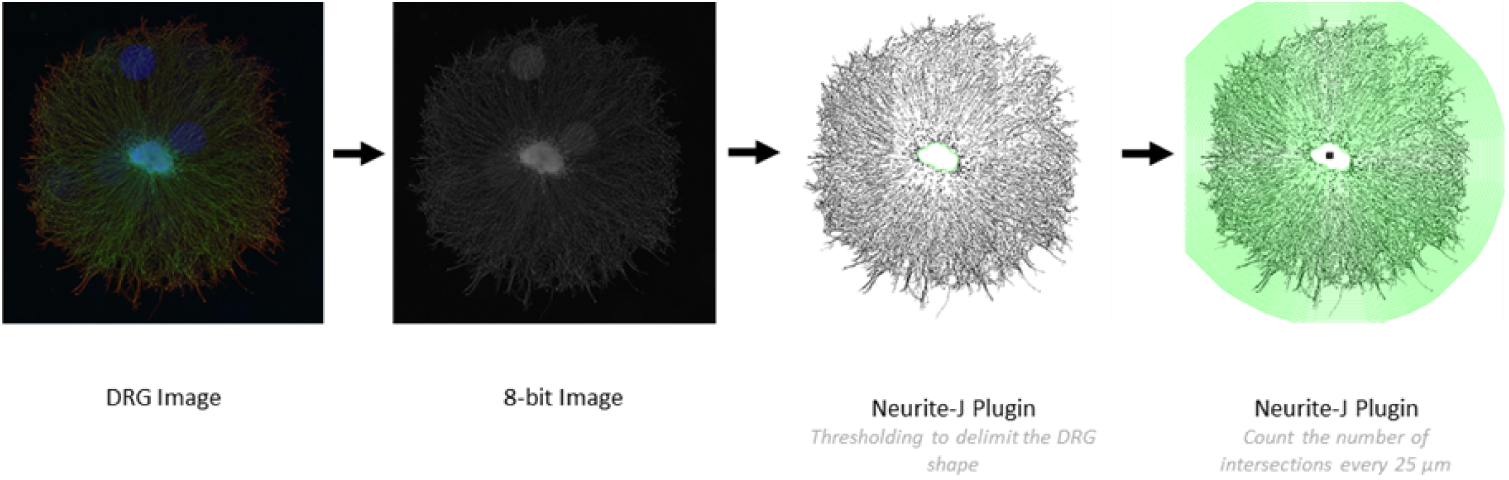
1mage treatment for axonal growth analysis. The image is first converted to an 8-bit format prior to being processed with Neurite-J plugin. The plugin counts the number of intersections with axons at 25 _µ_m intervals. The maximum distance is fixed when the number of intersections is below 10.

**Figure SI.8:**
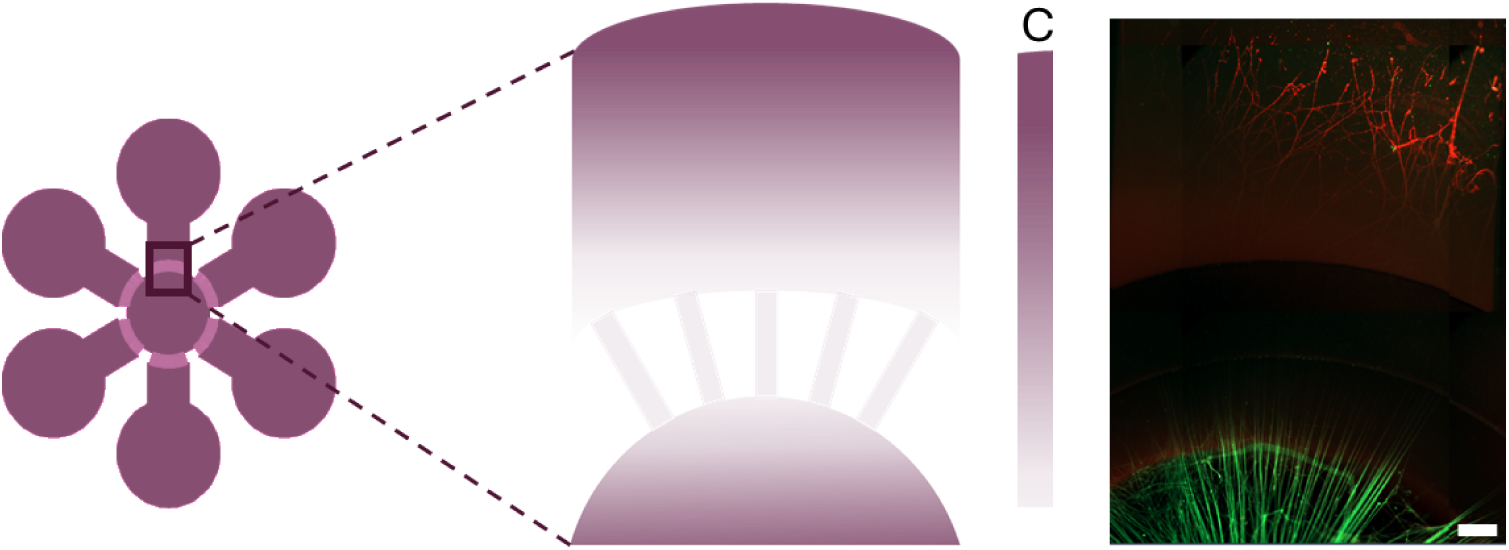
Schematic of the concentration gradient (C) close to the microchannels (right) the image was captured using an epifluorescence microscope and illustrate the difficulty of staining the axons. NFH (green) and SM1312 (red). Scale bar represents 100*µm*.

## 5 Manufacturing processes estimation

In Table SI.1, we detail the requirements of the SBC and PDMS manufacturing processes. In Table SI.2, we compare the environmental impacts of the products, from production to disposal, normalized against the impacts of PDMS. We compare it in a histogram - see Figure SI.10, 0.5 meaning the impact is equivalent for both polymer. A value *>* 0.5 indicates SBC is more impactful than PDMS and reciprocally.

**Figure SI.9:**
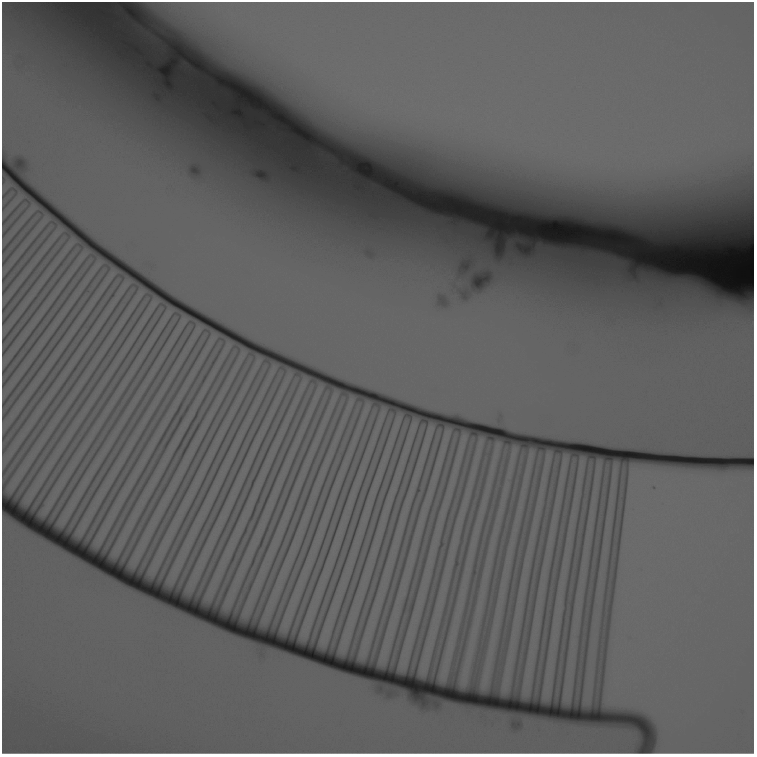
Brightfield image of the microchannels after four uses of the chip, indicating that they are no longer linked to the central chamber. Scale bar represents 100*µm*.

**Figure SI.10:**
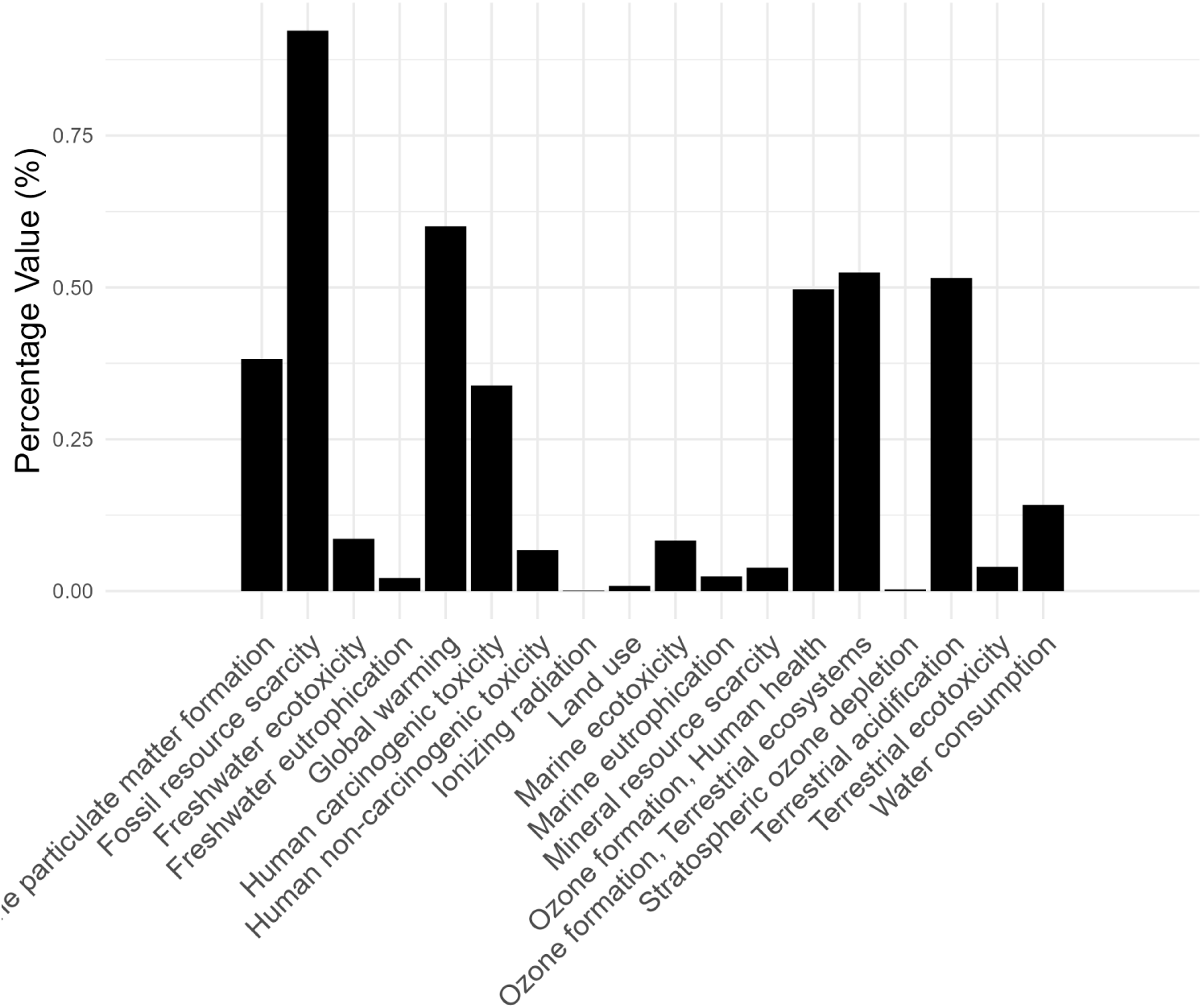
Ratio percentage of impact of SBC compared to PDMS.

**Figure SI.11:**
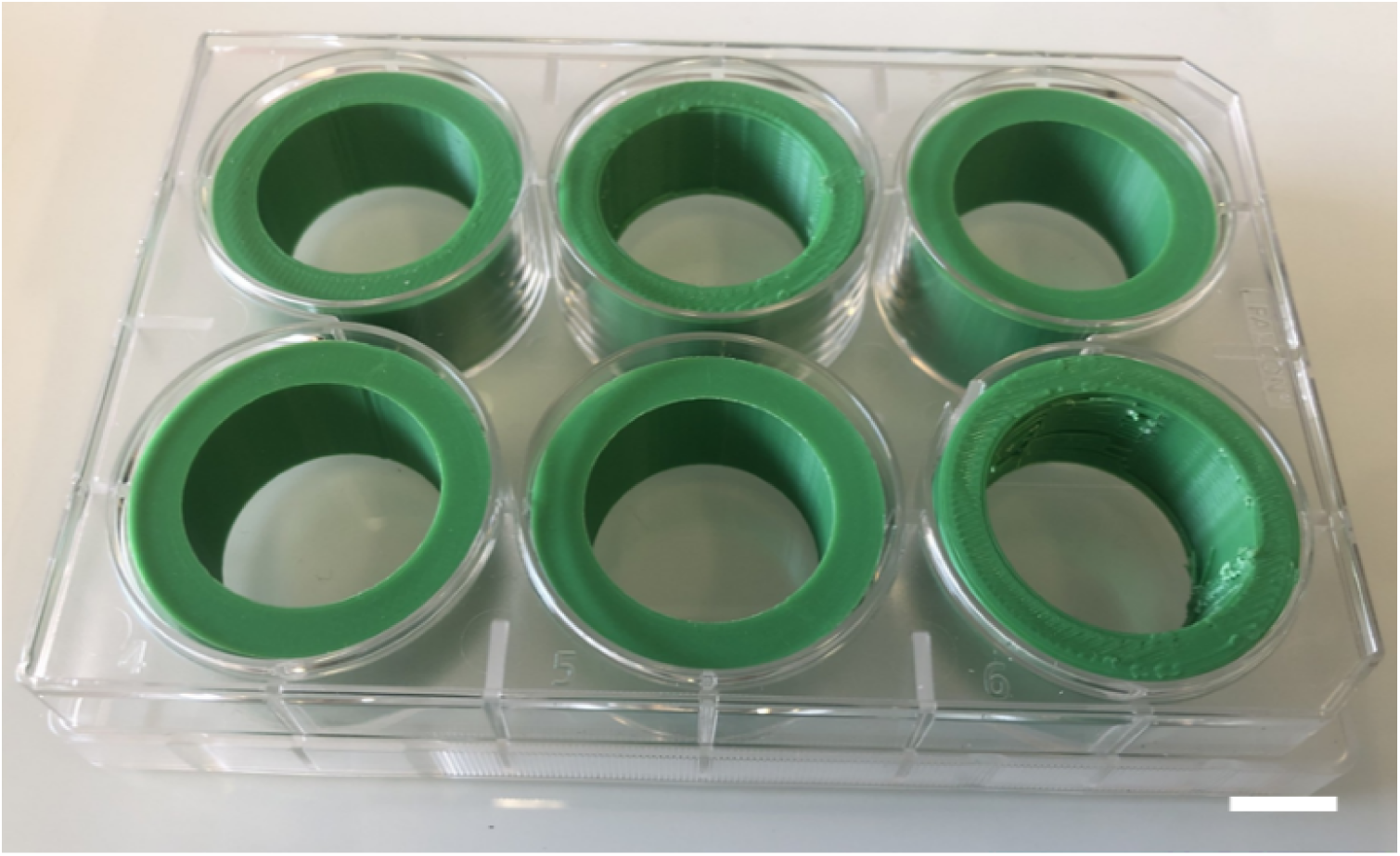
Picture of rings 3D printed and placed inside a 6-well plate to reduce the volume. Scale bar = 1*cm*.

**Figure SI.12:**
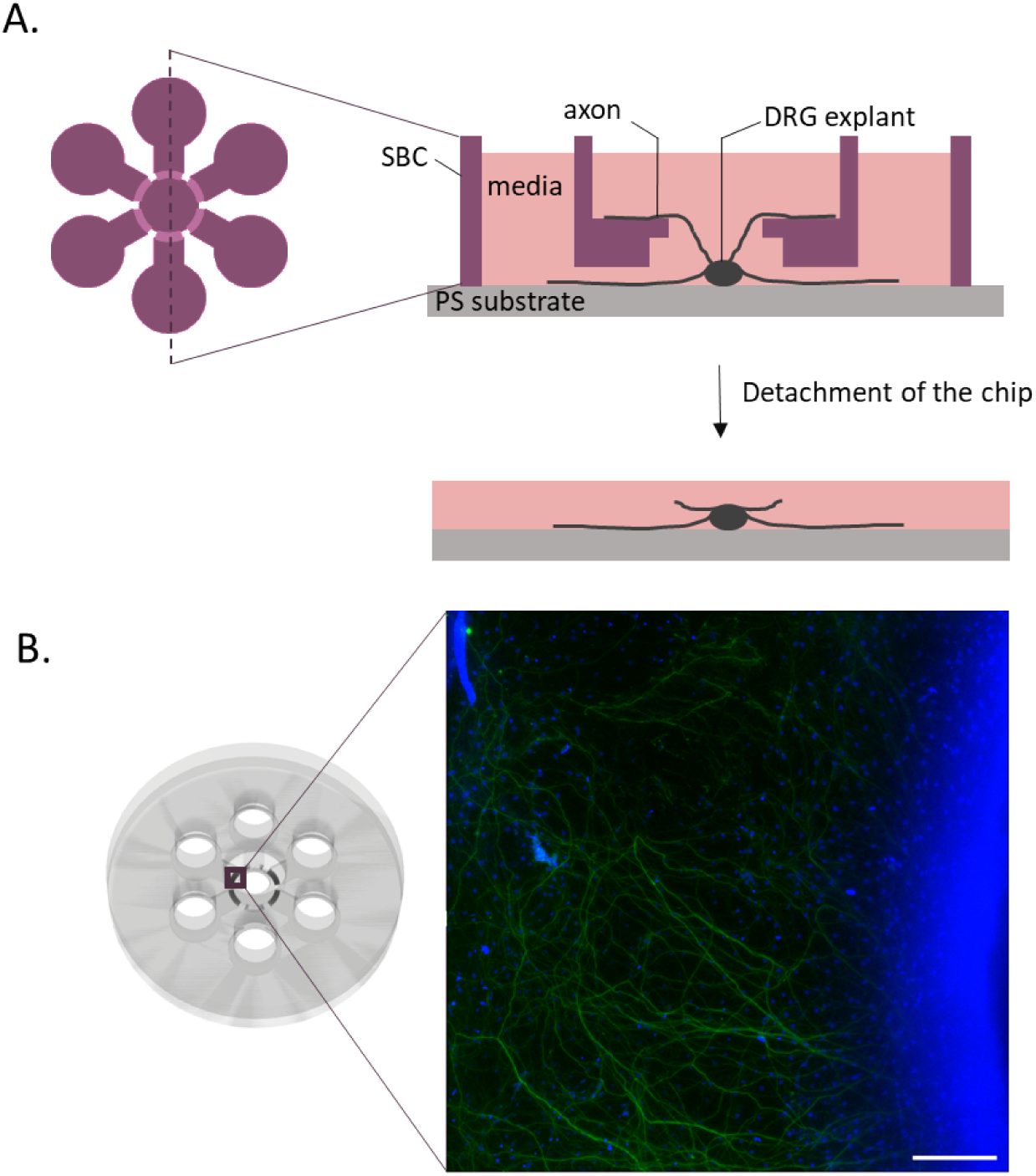
Scheme of the axons growing on top of the reservoir layer and the tearing of these axons when detaching the chip. B. 1mage captured using an epifluorescence microscope of the axons on top of the reservoir. NFH (green) and nuclei (NucRed). Scale bar represents 200*µm*.

## 6 Immunostaining volume optimization

Movie SI.1 shows the chip detachment with a fixed DRG at D1V15. Once detached, a green printed part is added to each well of the plates to minimize the volue used for immunostaining - see Figure SI.11.

The detachment of the chip tears the axons growing on top of the central reservoir - see Figure SI.12.

## 7 Protein and RNA extraction

The extraction of the proteins were attempted varying the number of DRG pulled after extraction from 4-well plates. The advantage of pulling 8 DRG is a reduced standard deviation of the sample, though the *DRG* = 1 condition displays a reasonable one. The proportion of SOX10 protein detected seems also reasonaby higher between 1 and 8, indicating that pulling strategies can be relevant for rare protein samples - see Figure SI13.

**Figure SI.13:**
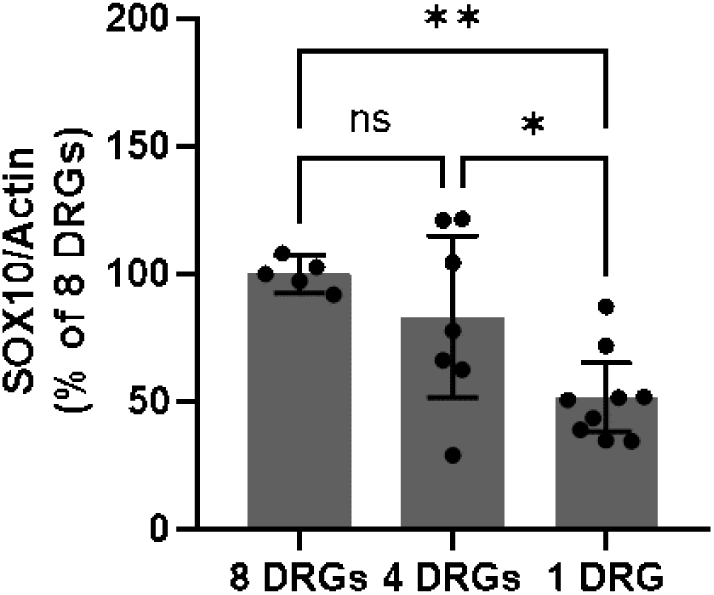
Comparison of Sox10 expression normalized by actin in 1, 4 and 8 DRGs cultivated for 11 days in 4-well plates. *β*-Actin was used as a loading control. Results represent the mean *±* SEM of three independent experiments (*n* = 5 *−* 9, embryos *e* = 3). A one-way ANOVA following by a Tukey’:s multiple comparisons test was performed for statistical analysis. The alpha value threshold was set at 5% and the p-values are represented as follows: ns = non-significant, \**p <* 0.05, \*\**p <* 0.01.

**Figure SI.14:**
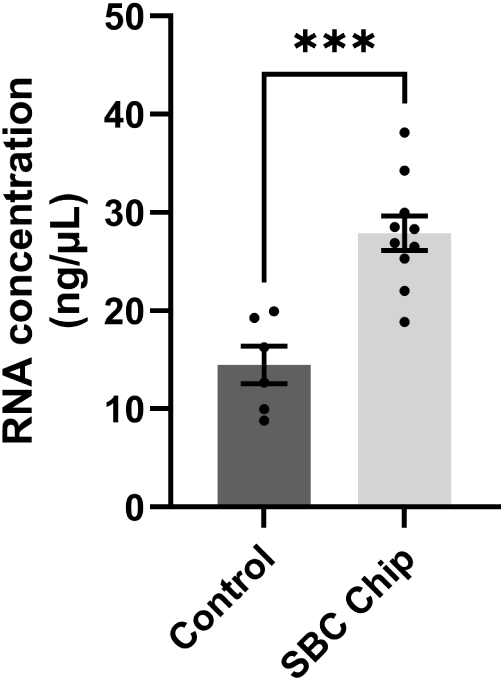
Comparison of Sox10 expression normalized by actin in 1, 4 and 8 DRGs cultivated for 11 days in 4-well plates. *β*-Actin was used as a loading control. Results represent the mean *±* SEM of three independent experiments (*n* = 5 *−* 9, embryos *e* = 3). A one-way ANOVA following by a Tukey’:s multiple comparisons test was performed for statistical analysis. The alpha value threshold was set at 5% and the p-values are represented as follows: ns = non-significant, \**p <* 0.05, \*\**p <* 0.01.

The comparison of the amount of RNA recovered in control against reversibly bonded chips revealed that the SBC chips provide more material in equivalent culture conditions - see Figure SI14. The quantification was performed on a Nanodrop (Thermo Fischer Scientific, USA).

